# Distributing task-related neural activity across a cortical network through task-independent connections

**DOI:** 10.1101/2022.06.17.496618

**Authors:** Christopher M. Kim, Arseny Finkelstein, Carson C. Chow, Karel Svoboda, Ran Darshan

## Abstract

Task-related neural activity is widespread across populations of neurons during goal-directed behaviors. However, little is known about the synaptic reorganization and circuit mechanisms that lead to broad activity changes. Here we trained a limited subset of neurons in a spiking network with strong synaptic interactions to reproduce the activity of neurons in the motor cortex during a decision-making task. We found that task-related activity, resembling the neural data, emerged across the network, even in the untrained neurons. Analysis of trained networks showed that strong untrained synapses, which were independent of the task and determined the dynamical state of the network, mediated the spread of task-related activity. Optogenetic perturbations suggest that the motor cortex is strongly-coupled, supporting the applicability of the mechanism to cortical networks. Our results reveal a cortical mechanism that facilitates distributed representations of task-variables by spreading the activity from a subset of plastic neurons to the entire network through task-independent strong synapses.

## Introduction

Large-scale measurements of neural activity show that learning can rapidly change the activity of many neurons, resulting in widespread changes in task-related neural activity [1, 2, 3, 4, 5]. For instance, a goal-directed behavior involving motor planning produces macroscopic neural dynamics that is coordinated by multiple cortical areas [1, 3], and neural correlates of action and choice made during a decision-making task are organized spatially across brain regions [4].

To gain insights into the circuit mechanism behind the observed widespread activity, it is critical to understand how interconnected neural circuits modulate their synaptic connections to produce the observed changes in task-related neural activity. Tracking synaptic modifications during learning [6, 7, 8, 9] and manipulating them to demonstrate a causal link with behavioral outputs [10, 11, 12, 13, 14], show that synaptic plasticity underlies learned behaviors and changes in neural activity [15, 16]. However, it is highly challenging to conduct multi-scale experiments that monitor and manipulate learning-specific synaptic changes at cellular resolution across a wide region of cortical networks, while measuring the resulting neural activity [17]. Thus, it remains unclear what aspects of the synaptic connections are modified to produce the widespread changes in task-related neural activity.

Here we investigated if task-related activity, learned locally in a dedicated subset of neurons, can spread across the network through pre-existing, task-independent, synaptic pathways. Although distributed neural activity may result from broad changes in synaptic connections across a neural network, we hypothesized that recruiting only a small subset of neurons is sufficient to generate the distributed task-related neural activity. To test this hypothesis, we used recurrent neural networks that provide a powerful data-driven approach for investigating how synaptic modifications can support the observed task-related neural activity. [18, 19, 20, 21, 22].

In typical implementations of network training, all the neurons in the network are considered to be plastic, in that the activity of every neuron is fit to the activity of experimentally recorded neurons, thereby constraining the entire network activity to the neural data [19, 20, 21]. In this study, we instead trained only a subset of neurons in a biologically plausible network to reproduce the activity of recorded neurons. The network consisted of excitatory (E) and inhibitory (I) populations of spiking neurons with sparse and strong connections [23, 24, 25]. Such pre-existing, task-independent, connections made the network settle into a cortical-like dynamical regime, where excitation and inhibition balanced each other [23, 25, 26], resulting in temporally irregular spikes and heterogeneous spike rates [27].

We applied our modeling framework to study the spread of task-related activity in the anterior lateral motor cortex (ALM) of mice performing a memory-guided decision-making task [21]. Similarly to neurons in the primate motor cortex [28, 29, 30], the activity of many neurons in ALM ramps slowly during motor preparation and is selective to future actions [21, 31, 32]. These task-related activity patterns are widely distributed across the ALM and are highly heterogeneous across neurons.

When a small number of synapses was trained to reproduce the ALM activity in a subset of neurons, we found that, surprisingly, the emerging activity in the untrained model neurons closely matched the responses of ALM neurons held out from training. In other words, the task-related ALM activity, learned in a subset of neurons, spread to other untrained neurons in the network without further training and produced activity that resembled the actual responses of ALM neurons. Analysis of the trained networks revealed that the pre-existing strong synapses between the neurons mediated the propagation of the task-related activity. The trained activity failed to spread in networks of neurons that were not strongly coupled. Optogenetic perturbation experiments of ALM activity provided additional evidence that the ALM network is strongly coupled, supporting the proposed mechanism for the spread of trained activity.

Our work provides a general circuit mechanism for spreading activity in cortical networks. It suggests that task-related activity observed in cortical regions during behavior can emerge from a subset of neurons with sparse synaptic reorganization and propagate to the rest of the network through the strong, task-independent synapses.

## Results

### Training strongly coupled spiking neural networks with sparse synapses

Our spiking network was based on a cortical circuit model that provided mechanistic explanation of the canonical features of cortical activity, such as temporally irregular spike trains, large trial-to-trial variability and a wide range of firing rates across neurons [23, 24, 25, 27, 33]. It consisted of excitatory (E) and inhibitory (I) neurons sparsely and randomly connected by strong synapses. This initial EI network structure, due to its random connectivity, was independent of the task to be learned. In addition, the strong excitatory and inhibitory synaptic inputs were dynamically balanced to maintain a stable network state, known as the *balanced regime*. As in the cortex, neurons, driven by fast fluctuating synaptic inputs, emitted spikes stochastically in this network state.

Here we describe a training scheme referred to as *Subset Training*, which trains a subset of neurons in the EI spiking network to generate target activity patterns, with the rest of the neurons untrained (**Fig. 1A, left**). After training the selected subset of neurons, we can analyze if the learning-related changes in activity spread throughout the network (**Fig. 1A, right**).

**Figure 1:**
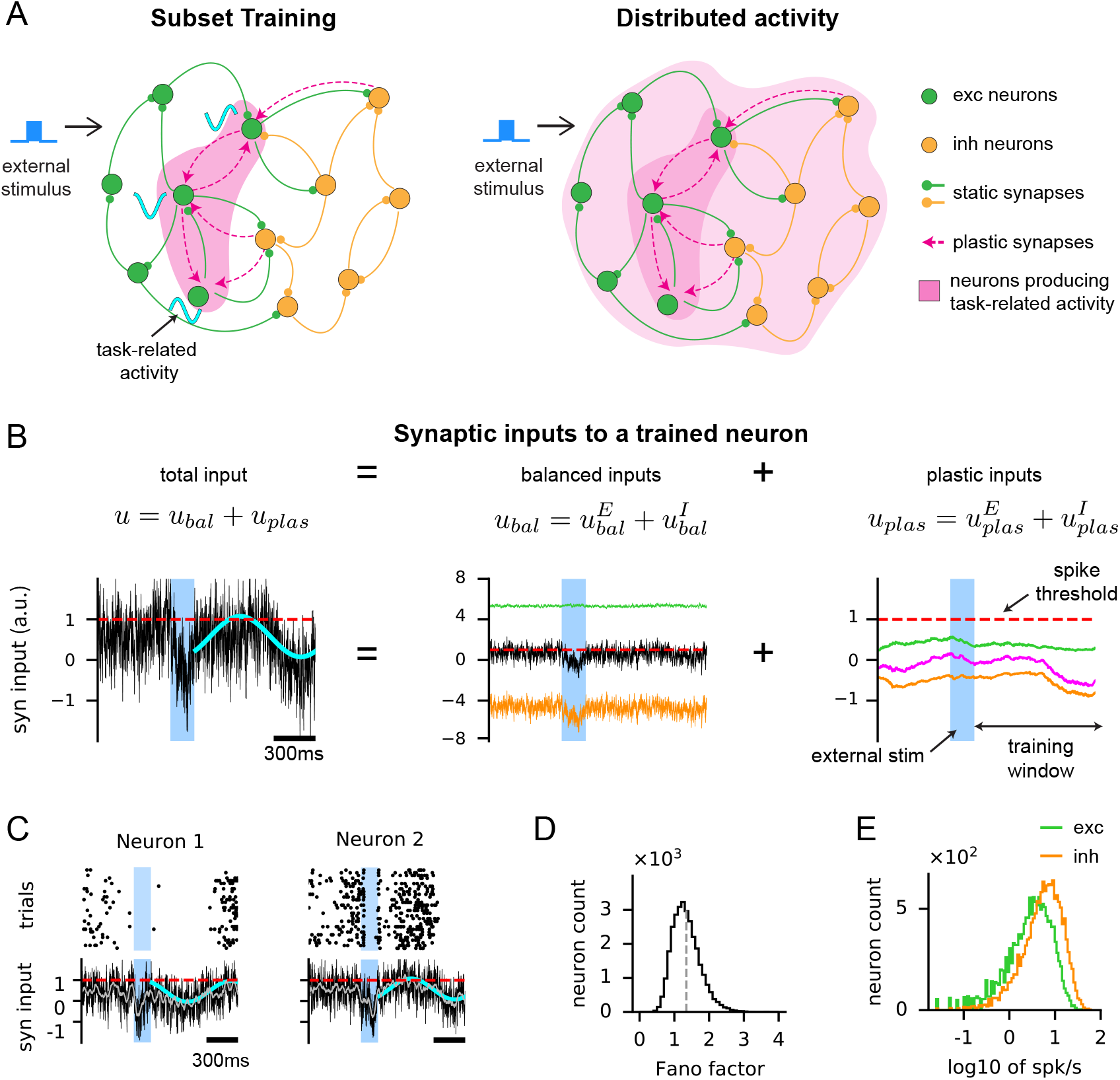
Training a subset of neurons in a strongly coupled spiking neural network with sparse plastic synapses. **(A)** Schematic of the Subset Training method (left). The network consisted of excitatory (green) and inhibitory (orange) neurons. Selected neurons (dark magenta, left) were trained to generate task-related activity patterns, modeled here as 1Hz sine waves with random phases (cyan curves), by modifying plastic synapses (dashed arrows, magenta) to the selected neurons. The static synapses (solid arrows, excitatory: green, inhibitory: orange) remained unchanged throughout training. External stimulus (blue pulse) triggered the neurons to generate the trained activity patterns. After training (right), task-related activity could potentially spread to the rest of the untrained neurons (light magenta, right) **(B)** The total synaptic input (left panel) to a trained neuron followed the target pattern (cyan) when triggered by an external stimulus (blue region). The total input is the sum of the balanced input, denoted as *u_bal_*, from static synapses (black; middle panel) and the plastic input, denoted as *u_plas_*, from plastic synapses (magenta; right panel). The balanced and plastic inputs can be further divided into excitatory (green) and inhibitory (orange) inputs. The spike-threshold of the neuron is at 1 (red dotted line). Note the scale difference between the balanced and plastic inputs. **(C)** Additional examples of the total synaptic inputs (same as the left panel in (B)) to trained neurons (bottom) following the target patterns (cyan); the 200ms moving average is shown in gray. Spike trains emitted by the neurons across 30 trials are shown on the top. **(D)** Fano factor of spike counts across 30 trials. **(E)** The log of firing rates of trained neurons. All neurons in the network were trained in this demonstration of the Subset Training method.

To model the effects of learning, a relatively small number of plastic synapses connecting to the subset of neurons was introduced to the existing EI spiking network. We implemented such network architecture by decomposing the recurrent connectivity into static and plastic components. The static component was the strongly coupled EI network connectivity that made the network operate in the balanced regime ( **Fig. 1A**, solid arrows). It remained unchanged throughout network training. For the plastic component, we added sparse connections, that were sparser than the EI connectivity (**Fig. 1A**, magenta dashed arrows). The choice of sparse plastic synapses was, in part, motivated by the synaptic connectivity found in the cortex that is sparse but functionally biased [34]. For instance, with *K* = 1000 static synapses, there were of the order of 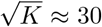 plastic synapses per neuron (**Table 2**). The plastic synapses were connected only to the trained neurons from randomly selected presynaptic neurons. During network training, a synaptic learning rule based on recursive least squares [19, 35, 36, 37] optimized the strength of plastic synapses to generate the target activity patterns in the selected subset of neurons.

**Table 1:** Default simulation and training parameters. Any differences from the above parameters are described in Table2.

| Neuron parameters |  | Values |
| --- | --- | --- |
| $\delta t$ | simulation time step | 0.1 ms |
| $\tau_m$ | membrane time constant | 10 ms |
| $v_{thr}$ | spike threshold | 1 |
| $v_{reset}$ | voltage reset after spike | 0 |
| Network parameters |  |  |
| $N$ | number of neurons | 30000 |
| $N_E$ | number of excitatory neurons | $N/2$ |
| $N_I$ | number of inhibitory neurons | $N/2$ |
| $p$ | connection probability | 0.2 |
| Synaptic parameters |  |  |
| $\tau_{bal}$ | static synaptic time constant | 3 ms |
| $\tau_{plas}$ | plastic synaptic time constant | 150 ms |
| $K$ | average number of static synapses to a neuron | $pN$ |
| $K_E$ | average number of excitatory static synapses to a neuron | $pN_E$ |
| $K_I$ | average number of inhibitory static synapses to a neuron | $pN_I$ |
| $L$ | number of plastic synapses to a neuron | see Table2 |
| $J_E$ | excitatory synaptic weight | $2.0/\sqrt{K_E}$ |
| $J_I$ | inhibitory synaptic weight | $-2.0/\sqrt{K_I}$ |
| $X$ | external input | $0.08\sqrt{K_I}$ |
| $J_{EE}$ | $E$ to $E$ static synaptic weight | $\gamma_E J_E$ |
| $J_{IE}$ | $E$ to $I$ static synaptic weight | $J_E$ |
| $J_{EI}$ | $I$ to $E$ static synaptic weight | $\gamma_I J_I$ |
| $J_{II}$ | $I$ to $I$ static synaptic weight | $J_I$ |
| $X_E$ | external input to excitatory neurons | $\gamma_X X$ |
| $X_I$ | external input to inhibitory neurons | $X$ |
| $\gamma_E$ | relative strength of $W_{EE}$ to $W_{IE}$ | 0.15 |
| $\gamma_I$ | relative strength of $W_{EI}$ to $W_{II}$ | 0.75 |
| $\gamma_X$ | relative strength of $X_E$ to $X_I$ | 1.5 |
| Training parameters |  |  |
| $\lambda$ | penalty for $L_2$-regularization | 0.05 |
| $\mu$ | penalty for ROWSUM-regularization | 8.0 |
| $N_{iter}$ | number of training iterations | 200 |
| $T_{target}$ | length of target patterns | 2 sec |

**Table 2:** The number of total neurons, trained neurons and plastic synapses in the simulated networks.

|  |  | Figure 2 | Figure 4 | Figures 1 & 5 |
| --- | --- | --- | --- | --- |
| Target functions |  | Neural PSTH | Synthetic PSTH | Sine function |
| Neurons |  |  |  |  |
| $N$ | # neurons | $5 \cdot 10^3$ | $3 \cdot 10^4$ | $3 \cdot 10^4$ |
| $N_{trained}$ | # trained neurons | 1824 | $3 \cdot 10^3$ to $1.5 \cdot 10^4$ | $3 \cdot 10^4$ & $1.5 \cdot 10^4$ |
| Static synapses to a neuron |  |  |  |  |
| $p$ | conn prob of static synapses | 0.2 | 0.2 | 0.213 |
| $K = pN$ | # static synapses to a neuron | **1,000** | **6,000** | **6,400** |
| Plastic synaptic weights to a trained neuron |  |  |  |  |
| $J_E, J_I$ | see Table 1 | | | |
| $W_{EE}$ | E to E plastic synaptic weight | $0.66J_E$ | $J_E$ | $2J_E$ |
| $W_{IE}$ | E to I plastic synaptic weight | $0.66J_E$ | $J_E$ | $2J_E$ |
| $W_{EI}$ | I to E plastic synaptic weight | $0.33J_I$ | $0.5J_I$ | $J_I$ |
| $W_{II}$ | I to I plastic synaptic weight | $0.33J_I$ | $0.5J_I$ | $J_I$ |
| Number of plastic synapses to a trained neuron |  |  |  |  |
| $\sqrt{K}$ | order of # plastic synapses | 32 | 77 | 80 |
| $L_{rec} = c\sqrt{K}$ | # recurrent plastic synapses | 264 | 440 | 226 |
| $L_{ffwd} = c\sqrt{K}$ | # ffwd plastic synapses | 300 | 200 | 0 |
| $L = L_{rec} + L_{ffwd}$ | # total plastic synapses | **564** | **640** | **226** |
| Sparsity of plastic synapses |  |  |  |  |
| $L/K$ | # plastic / # static synapses | **0.564** | **0.106** | **0.035** |

This arrangement of plastic synapses, which connected only to the selected subset of neurons, allowed us to examine the role of the pre-existing recurrent connections of the EI network in spreading the trained activity to the untrained neurons, which were not targeted by the plastic synapses. In addition, due to their sparsity, the plastic synaptic inputs were substantially weaker than the strong excitatory and inhibitory synaptic inputs of the existing EI network (**Fig. 1B**). This allowed the network to stay in the balanced regime after training and supported robust network training, independent of the density of synaptic connections (**Fig. S1**; see Methods for full description of the training and details on the sparse plastic synapses).

In the trained network, the total synaptic input to each trained neuron was able to successfully follow the target patterns (**Fig. 1B, left**; **Fig. 1C**). The statistics of spiking activity of the trained network were similar to those of untrained, strongly coupled EI networks, thus consistent with the spiking activity of cortical neurons. Specifically, due to the highly fluctuating balanced input, the spike trains of each neuron were irregular and exhibited large trial-to-trial variability (**Fig. 1D**, Fano factor = 1.4) [23, 24, 38]. The firing rate distribution was also highly skewed and was well approximated by a log-normal distribution (trained model: **Fig. 1E**, neural data: **Fig. S2D**) [27].

### Spread of trained neural activity to untrained neurons

We applied the Subset Training method to reproduce the firing rate patterns of cortical neurons recorded from the anterior lateral motor cortex (ALM) during a memory-guided decision-making task [21]. Mice learned to respond to an optogenetic stimulation of neurons in the vibrissal somatosensory cortex (vS1) by licking right when stimulated and licking left otherwise (**Fig. 2A**). For training networks and analysis of trained networks, we used the electrophysiological recordings in ALM of the spiking activity of putative pyramidal neurons (excitatory; *N_pyr_* = 1824) and putative fast-spiking neurons (inhibitory; *N_fs_* = 306) when the mice responded correctly to lick-right and lick-left conditions.

**Figure 2:**
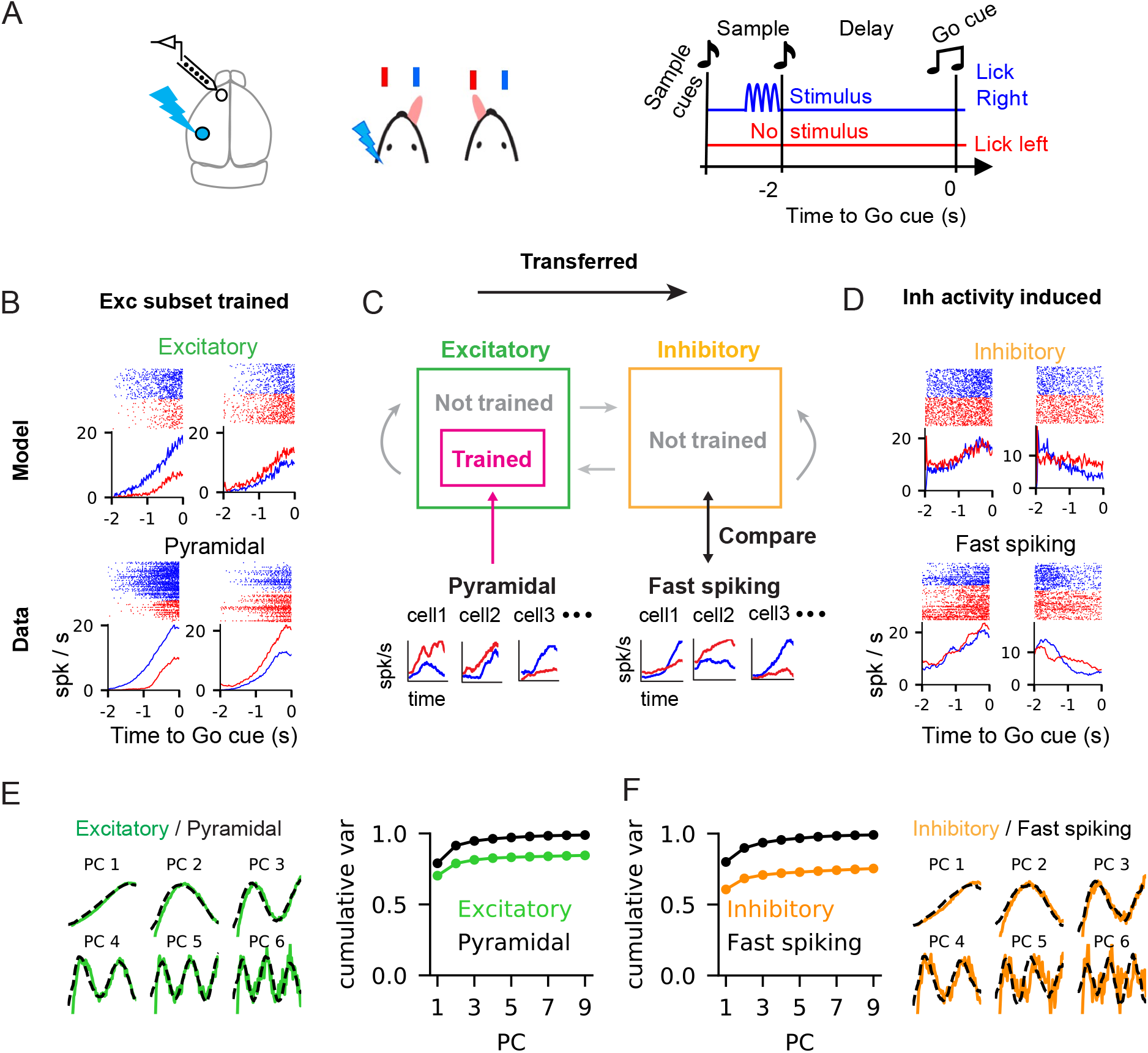
Reproducing ALM activity in a subset of neurons and the spread of trained activity to the entire network. **(A)** Schematic of experimental setup. Mice learned to lick right when optogenetic simulation was delivered to somatosensory neurons and to lick left when there was no stimulation. The spiking activity of ALM neurons was recorded during the task. **(B)** Trial-averaged firing rates and raster plots of the spike trains across multiple trials (lick-right: blue, lick-left: red). Trained excitatory model neurons (top) and pyramidal ALM neurons used for training the excitatory model neurons (bottom). **(C)** A subset of excitatory neurons in the spiking neural network learned to reproduce the PSTHs of pyramidal ALM neurons. The rest of the neurons in the network were not trained. After training, the activity of untrained inhibitory model neurons was compared to the activity of fast-spiking ALM neurons. **(D)** Trial-averaged firing rates and raster plots of the spike trains across multiple trials (lick-right: blue, lick-left: red). Untrained inhibitory model neurons (top) and fast-spiking ALM neurons (bottom) that best resembled the PSTHs of the inhibitory model neurons (see also Fig. 3 and Fig. S4). **(E, F)** The principal components (PCs) of the PSTHs of excitatory/pyramidal and inhibitory/fast-spiking neurons (model/data) and the cumulative variances explained by the PCs.

We asked what aspects of the network connectivity should change to reproduce the activity of ALM neurons in a strongly coupled spiking neural network. In previous studies, in which networks were trained to generate specific patterns of neural activity, all the units in the network were trained to reproduce the target activity patterns [18, 19, 20, 21]. Here, we trained only a *subset* of neurons, embedded in the strongly coupled EI network, to reproduce the spiking activity of ALM neurons. By analyzing the activity of neurons in the trained network, we found that synaptic reorganization to a subset of neurons was sufficient to generate the observed ALM activity throughout the entire network, including the *untrained* neurons. Importantly, the spread of target activity patterns from the subset of trained neurons to the rest of neurons was not observed in a network that was not strongly coupled (**Fig. S9**). This suggests that the spread of trained activity to untrained neurons is a characteristic of strongly-coupled networks, but not a general outcome of recurrent networks.

The training targets were the trial-averaged peri-stimulus time histogram (PSTH) of pyramidal neurons recorded in ALM during the delay period (**Fig. 2B, bottom**). We trained a subset of excitatory neurons in the model network to reproduce the target activity patterns (**Fig. 2C**). Each trained neuron received recurrent plastic synapses from randomly selected excitatory and inhibitory neurons in the network and feedforward plastic synapses from external neurons, which accounted for the potential inputs from outside the ALM. By modifying the plastic synapses, the trained neurons reproduced two PSTHs, corresponding to lick-right and lick-left conditions, when evoked by two different stimuli. The rest of the excitatory, as well as all of the inhibitory neurons in the network, were not trained (**Fig. 2C**).

After training, the firing rate of trained excitatory neurons successfully reproduced the PSTHs of pyramidal neurons (**Fig. 2B, top; Fig. S2A,B**), even though the plastic synaptic inputs were substantially weaker than the excitatory and inhibitory inputs from the static synapses (**Fig. S2C, right**). We estimated the correlations between single neuron PSTHs in the model and in the data (**Fig. S2C, left**), as well as the similarity in their population activity (**Fig. 2E, left**) to asses the success of the training. For the latter, we performed Principal Component Analysis (PCA) on the PSTHs of neurons, which is a dimensionality reduction technique used for identifying a set of activity patterns that captures a large fraction of variance in the population activity. The principal components (PCs) of the PSTHs of the trained excitatory neurons closely matched the PCs of the pyramidal neurons. Moreover, the first six PCs explained close to 80% of the trained neurons’ activity, thus the recurrent network displayed low-dimensional dynamics as in the pyramidal neurons in ALM (**Fig. 2E, right**) [39].

Next, we examined the activity of the *untrained* neurons. Similarly to the trained excitatory neurons, their activity tended to ramp before go-cue and was choice-selective (**Fig. 2D, top**). The PCs of their PSTHs were almost identical to the trained excitatory neurons (**Fig. 2F, right**; **Fig. S3E**). This finding showed that cortical-like activity generated within the subset of excitatory neurons spread to the the rest of the network without additional synaptic reorganization to the untrained neurons.

Finally, we found that the PCs of the PSTHs of the fast-spiking ALM neurons, whose activity was not learned by the network, were almost identical to the PCs of the untrained inhibitory model neurons (**Fig. 2F, right**). For the lick-left trial, the higher PC modes (i.e., PC4 and PC5) of the inhibitory model neurons and fast-spiking ALM neurons (**Fig. S3E, left**) both mildly deviated from the corresponding PCs of trained neurons (**Fig. S3C, left**), but still closely resembled each other. A good agreement between the untrained model neurons and the held-out neural data supported the hypothesis that cortical-like activity learned within a subset of neurons can spread and produce cortical-like activity in the entire network. This could explain why the activity of putative fast-spiking neurons in ALM is heterogeneous, yet is very similar to the activity of putative pyramidal neurons [21].

### Similarity in the neural activity of untrained model neurons and ALM neurons

To further investigate the similarities between the activities of the untrained inhibitory neurons in the trained network and the fast-spiking ALM neurons, we compared their PSTHs at the single neuron and population levels.

At the single neuron level, we identified an untrained inhibitory neuron that best matched each fast-spiking ALM neuron, based on the mean-squared-error of the PSTHs of all possible pairings between the ALM neuron and the population of inhibitory model neurons. Figure 3A shows the PSTHs of several matched pairs and their correlations for the lick-right and lick-left trials (see **Fig. S4** for all the matched pairs). Evaluating the correlations of all the matched pairs showed that they were significantly higher than the spurious correlations obtained by matching the fast-spiking ALM neurons to inhibitory neurons in an untrained balanced network (**Fig. 3B, left**; two sample Kolmogorov-Smirnov tests; p-value < 0.0001).

**Figure 3:**
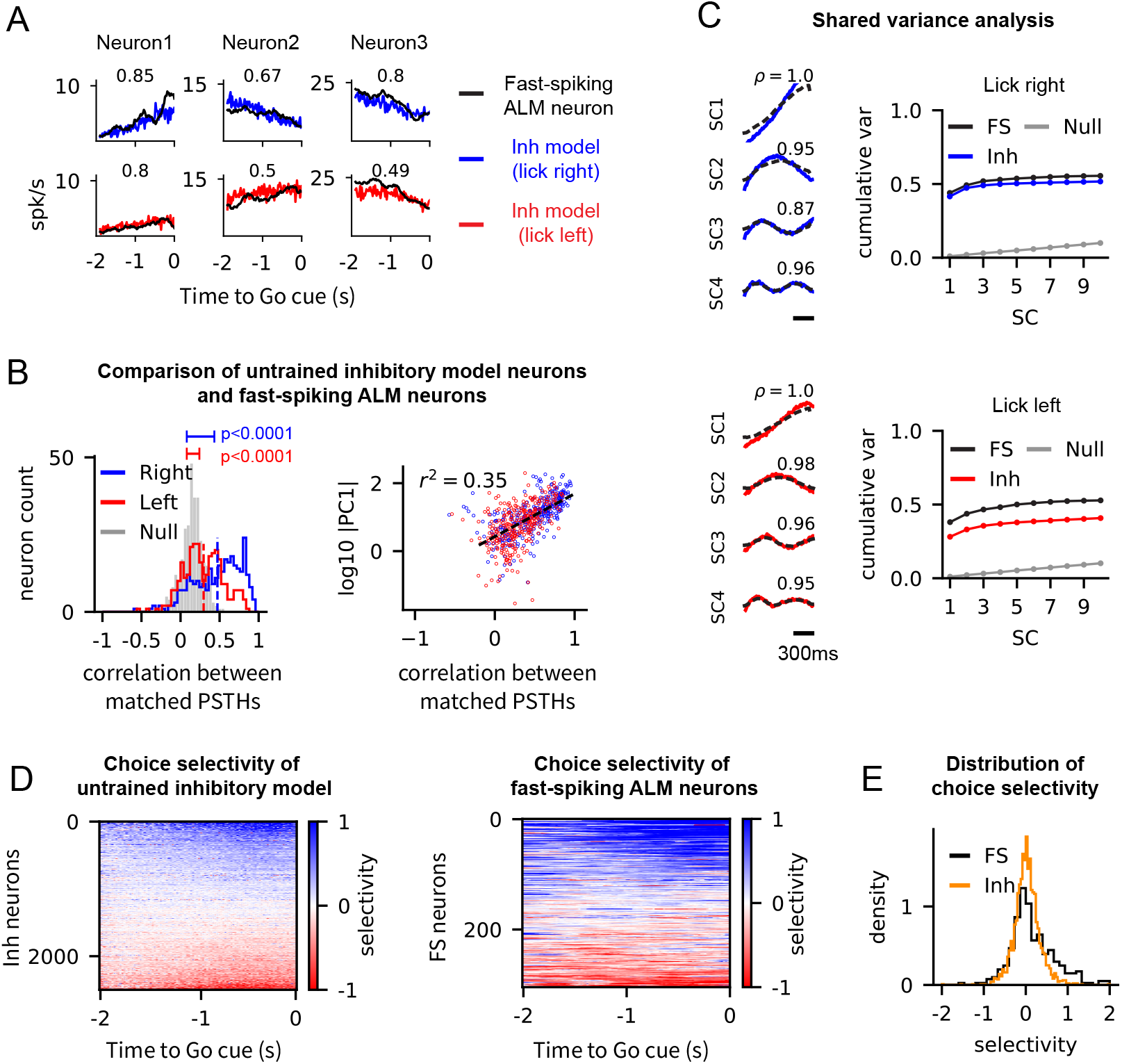
Untrained inhibitory neurons in the trained network display similar task-related activity to the fast-spiking neurons recorded in the ALM. **(A)** Examples of the PSTHs of untrained inhibitory neurons (lick-left: red, lick-right: blue) that best fit the PSTHs of fast-spiking ALM neurons (black). Correlations of the matched pairs are shown in each panel. **(B)** Correlations between the PSTHs of all the matched inhibitory model neurons and the fast-spiking ALM neurons for the lick-right and lick-left trial types (left). The null network shows the correlation between the PSTHs of the fast-spiking ALM neurons and the best-fit neurons in the initial balanced network, i.e., without training. The PSTHs of the neurons in the trained and the null networks were both obtained by averaging the spike trains from 400 trials starting at random initial conditions. The p-values of the Kolmogorov-Smirnov tests between the trained and null networks for both trial types are shown. PC1 (right) represents the projection of the PSTH of a fast-spiking ALM neuron onto the first PC, i.e., the ramping mode (see Fig. 2F). R-squared value of the linear regression is shown. **(C)** Shared components (SC) and the cumulative shared variance explained by them for the lick-right (top) and lick-left trial types (bottom). The null network shows the shared variance between the fast-spiking ALM neurons and the initial balanced network. **(D)** Choice selectivity of all the untrained inhibitory neurons in the trained network (left) and the fast-spiking ALM neurons (middle). Choice selectivity was defined at each time point as the difference of the PSTHs to the lick-right and lick-left trial types, and then normalized by the average firing rate of each neuron. **(E)** Distribution of choice selectivity of untrained inhibitory neurons in the trained network (orange) and fast-spiking ALM neurons (black). Choice selectivity of a neuron shown here was obtained by averaging the choice selectivity over the 2 second time window shown in (D).

To elucidate which aspects of the fast-spiking ALM activity were captured by the untrained inhibitory neurons in the trained network, we examined if certain activity patterns of the fast-spiking ALM neurons were indicative of the goodness-of-fit to the model neurons. We found that the projection of the PSTHs of the fast-spiking ALM neurons onto their first PC, a ramping mode that captured over 70% of the variance in the ALM activity (see PC1 in **Fig. 2F**), was a good indicator for how well the untrained model neurons could fit the fast-spiking ALM neurons (**Fig. 3B, right**). This analysis suggested that the ramping mode was the dominant component of the trained activity that was transferred to the untrained inhibitory neurons and shared with the fast-spiking ALM neurons.

We systematically examined the activity patterns shared by the populations of untrained inhibitory neurons and fast-spiking ALM neurons, by analyzing the shared-variance between the two population activities. The shared-variance analysis identified population vectors along which two population activities co-varied maximally and yielded population-averaged activity along those directions (shared components) and the proportion of variance explained by the shared components (shared variance; see [40] and **Methods** for details). The shared components (SCs) were similar to the PCs of the untrained inhibitory subnetwork and fast-spiking ALM activities (compare the SCs in **Fig. 3C** to the PCs in **Fig. 2F**), and the first four components captured most of the shared variance (**Fig. 3C**). In particular, consistent with the single neuron analysis shown in Fig. 3B, the first shared component was a ramping mode (SC1 in **Fig. 3C**).

In addition to the spiking activity patterns, we asked if functional properties, such as choice selectivity, were transferred to the untrained neurons in the trained network. It has been shown that pyramidal ALM neurons in mice display selectivity to the animal’s choice [21, 39, 41]. As expected, the excitatory model neurons, trained to reproduce the activity of pyramidal ALM neurons, also displayed choice selectivity (**Fig. S5**). Interestingly, we also found that the fast-spiking ALM neurons in the neural data were choice selective (**Fig. 3D,E**; see also Supplementary Fig. 2 in [21]). These observations led us to examine if the untrained inhibitory neurons in the trained network exhibited choice selectivity, as in the fast-spiking ALM neurons.

To this end, we analyzed the difference of the PSTHs to two trial types (lick-right versus lick-left) in all the untrained inhibitory neurons and found that they displayed choice selectivity (**Fig. 3D**). Moreover, the distribution of the choice selectivity of fast-spiking ALM neurons and untrained inhibitory neurons were in good agreement (**Fig. 3E**). This finding shows that not only the trained neural activity can propagate throughout the network, but the choice selectivity emerged in a subset of neurons can spread to the untrained parts of the network as well. In particular, this suggests an alternative mechanism for how selectivity may emerge in inhibitory neurons. In contrast to previous models that required specific connectivity between excitatory-inhibitory neurons for selective responses to emerge [42, 43], our model suggests that choice selectivity in inhibitory neurons can arise in strongly coupled networks even when the connections to the inhibitory neurons are non-specific.

### Spreading of trained neural activity improves if the inhibitory subnetwork is trained

So far, we showed that the cortical-like activity originating from the excitatory neurons can spread to the untrained inhibitory neurons. Next, we asked how the spreading of trained activity may depend on the type of neurons being trained. To address this question, we considered two training scenarios where either the excitatory or the inhibitory subnetwork (but not both) was trained to generate the target activity patterns (**Fig. 4A, right**).

**Figure 4:**
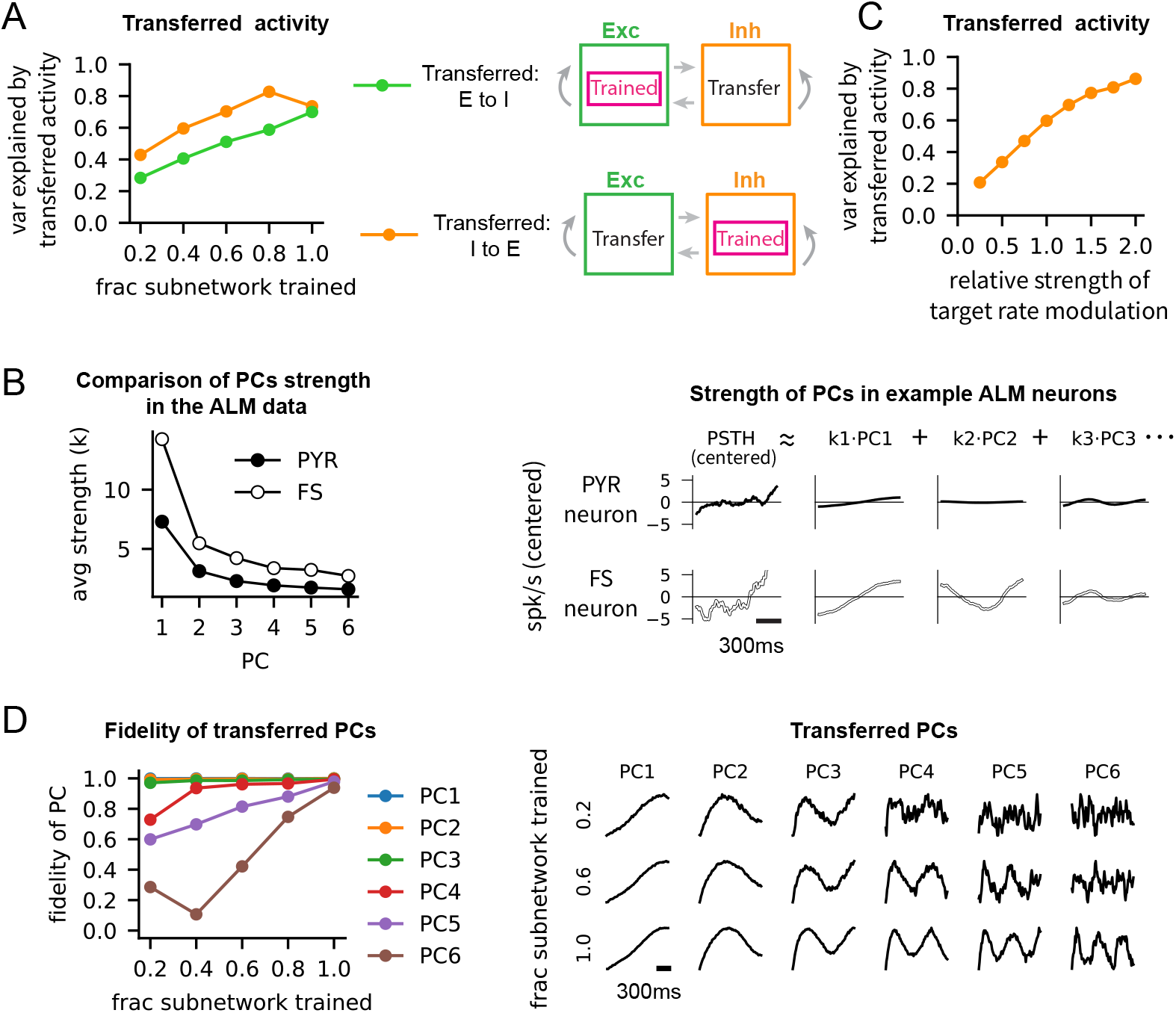
Trained activity originating from the inhibitory subnetwork spreads more effectively than the trained activity from the excitatory subnetwork. **(A)** Schematic of two training scenarios (right). A subset of neurons in the excitatory subnetwork (top) or the inhibitory subnetwork (bottom) was trained to reproduce synthetic neural activity. The fraction equals 1 (left) if all the neurons in the trained subnetwork are trained. The transferred activity was defined as the variance explained by the first six PCs of the PSTHs of all the neurons in the untrained subnetwork. **(B)** The strength of PCs of the PSTHs of pyramidal and fast-spiking ALM neurons (left). The absolute value of the loading of each PC on all the neurons in the population was averaged to obtain the average strength of each PC, denoted as *k*. Examples of centered PSTHs (right), i.e., mean rate subtracted, showing that the strength of the *i^th^* PC, denoted as *k_i_*, was stronger in the fast-spiking ALM neurons. **(C)** The modulation of the trained synthetic inhibitory rate was adjusted by scaling the centered PSTH by a multiplicative factor, referred to as the relative strength of modulation. For instance, it equals 2 if the centered PSTHs are doubled. As in (A), the transferred activity was defined as the variance explained by the first six PCs of the PSTHs of all the untrained excitatory neurons. The fraction of trained inhibitory neurons in the inhibitory subnetwork was 0.4. **(D)** The fidelity of transferred PCs (left) was defined by the correlation between the PCs of the trained and transferred activity. A subset of neurons in the excitatory subnetwork was trained, and the activity of the untrained inhibitory subnetwork was analyzed to obtain the transferred PCs. Examples of transferred PCs in the untrained inhibitory neurons (right) are shown, as the fraction of trained neurons is varied.

The number of fast-spiking ALM neurons recorded from the mice (*N_fs_* = 306) was, however, too small to train the inhibitory neurons in large-scale spiking neural networks. We thus developed a method to generate synthetic neural activity that had similar low-dimensional dynamics as the ALM neurons. Briefly, we first performed principal component analysis on the PSTHs of ALM neurons to obtain the PCs (**Fig. 2E, F**) and the empirical distribution of each PC’s loading onto the neurons. To construct a synthetic target activity for neuron *i*, we sampled (1) a baseline rate *r_i_* from the firing rate distribution and (2) each PC’s loading on the neuron from the empirical distribution, conditioned on the rate *r_i_* (see **Fig. S6** and Methods for details). Applying this method to the lick-left and lick-right trial types and to pyramidal and fast-spiking neuronal types, we were able to generate unlimited number of cortical-like PSTHs needed for training large-scale networks consisting of, e.g., *N* = 30, 000 neurons. In particular, these synthetic neural activity had statistically identical low-dimensional dynamics as the ALM neurons (**Fig. S6E**)

Using the synthetic neural activity as the target activity patterns, we performed the two training scenarios where we trained a subset of neurons in the excitatory or the inhibitory subnetwork to reproduce the synthetic neural activity. Following training, we compared the spiking activities of the untrained neurons in the subnetworks that were not trained.

We first observed that the PCs of synthetic neural activity was transferred to the untrained neurons when a sufficient number of neurons were trained (**Fig. 4D, right**). Such transfer of PCs was similar to what we found in the untrained inhibitory neurons when the excitatory neurons were trained on the activity of ALM pyramidal neurons (**Fig. 2E,F**). Based on the transfer of PCs and the low dimensionality of ALM activity, we used the variance explained by the first six PCs of the PSTHs of the untrained neurons to quantify the transferred cortical-like activity. In the trained neurons, the first six PCs explained 80% of the activity, regardless of the trained neuronal type (E or I) or the fraction of trained neurons (**Fig. S7A**). On the other hand, the cortical-like activity transferred to the untrained neurons gradually increased with the fraction of trained neurons. Moreover, the transferred activity was stronger by 20% when the inhibitory subnetwork was trained, compared to when the excitatory subnetwork was trained (**Fig. 4A, left**).

To understand what allowed the activity patterns of inhibitory neurons to spread better to the untrained neurons, we examined the differences in the spiking activities of the pyramidal and fast-spiking ALM neurons. The mean firing rate of each neuron was subtracted from its PSTH to remove the differences in the baseline firing rates of the pyramidal and fast-spiking ALM neurons (**Fig. 4B, right**). The principal component analysis of the centered PSTHs revealed that the strength of every PC was stronger in the fast-spiking neurons than in the pyramidal neurons, when the loadings on each PC were averaged over the population of neurons (**Fig. 4B, left**). This analysis showed that the modulation of firing rate around the mean rate was larger in the fast-spiking neurons, raising the possibility that stronger rate modulation leads to stronger activity transfer.

To test if stronger modulations in the trained activity patterns would increase the transferred activity to the untrained neurons, we adjusted the modulation strength of the synthetic inhibitory activity and trained a fixed subset of inhibitory neurons to generate target activity patterns with different levels of rate modulations. We found that, in the untrained excitatory neurons, the variance explained by the cortical-like activity increased monotonically with the modulation strength of the trained inhibitory neurons. (**Fig. 4C**). These results suggested that the stronger rate modulations in the fast-spiking ALM neurons enabled the trained inhibitory neurons in the model to spread their activity patterns to the untrained neurons more effectively. It also suggested that inhibitory neurons, whose baseline spiking rates are typically higher than the excitatory neurons in cortex (e.g., mean firing rates of ALM pyramidal and fast-spiking neurons were ~ 4Hz and ~ 11Hz, respectively, in our data), can support stronger rate modulations and potentially play a more significant role in spreading the trained activity patterns.

The finding that activity patterns with strong rate modulation spread better was also observed across the PCs. The lower PC modes of the ALM spiking activity showed stronger modulation than the higher PC modes, as expected, since the lower PC modes capture more variance (**Fig. 4B**). To quantify how well the trained PCs transferred to the untrained neurons, we computed the correlation between the PC modes of the trained and transferred activity (**Fig. 4D, left**). The lower PC modes (PC1 to PC3) transferred with high fidelity even when only 20% of the neurons were trained. On the other hand, the transfer of the higher PC modes (PC4 to PC6) improved gradually when the fraction of trained neurons increased. This result suggested that the lower PC modes, due to their strong modulations, can spread more robustly to the rest of the neurons, promoting low-dimensional neural dynamics across a strongly coupled network.

Taken together, our results demonstrate that trained activity patterns with stronger rate modulations, which can emerge from the fast-spiking ALM neurons or lower PC modes, have greater influence on the untrained neurons in the network.

### Network mechanism for distributing trained neural activity in strongly coupled networks

In a recurrent neural network, in which neurons are highly inter-connected, it may seem obvious that task-related activity can spread from one part of the network to another part through the connections that are not optimized for the task. However, this intuition becomes less clear when the activity of a neuron is determined by integrating a large number of heterogeneous presynaptic activities, as considered in our network model and is the case in the cortical network.

A close examination of networks with a large number of connections reveals that whether the task-related activity can spread depends on the operating regime of the network. If the synaptic connections to an untrained neuron randomly sample and sum a large number of heterogeneous activity patterns, one could expect that the activity patterns will be averaged-out at the level of the synaptic input. Then, the untrained neuron will not display any activity patterns, as shown in **Fig. S9**. In this section, we give an intuitive explanation that this is not the case if the network is strongly coupled and operates in the balanced regime (**Fig. 5**). In this regime, heterogeneities of presynaptic activities can be preserved in the synaptic inputs to untrained neurons because of the strong synapses and then manifested in the post-synaptic rate of the untrained neurons due to the dynamic cancellation of the large, unmodulated components of the excitatory and inhibitory inputs. A detailed explanation, together with a mathematical analysis, is given in the Methods.

**Figure 5:**
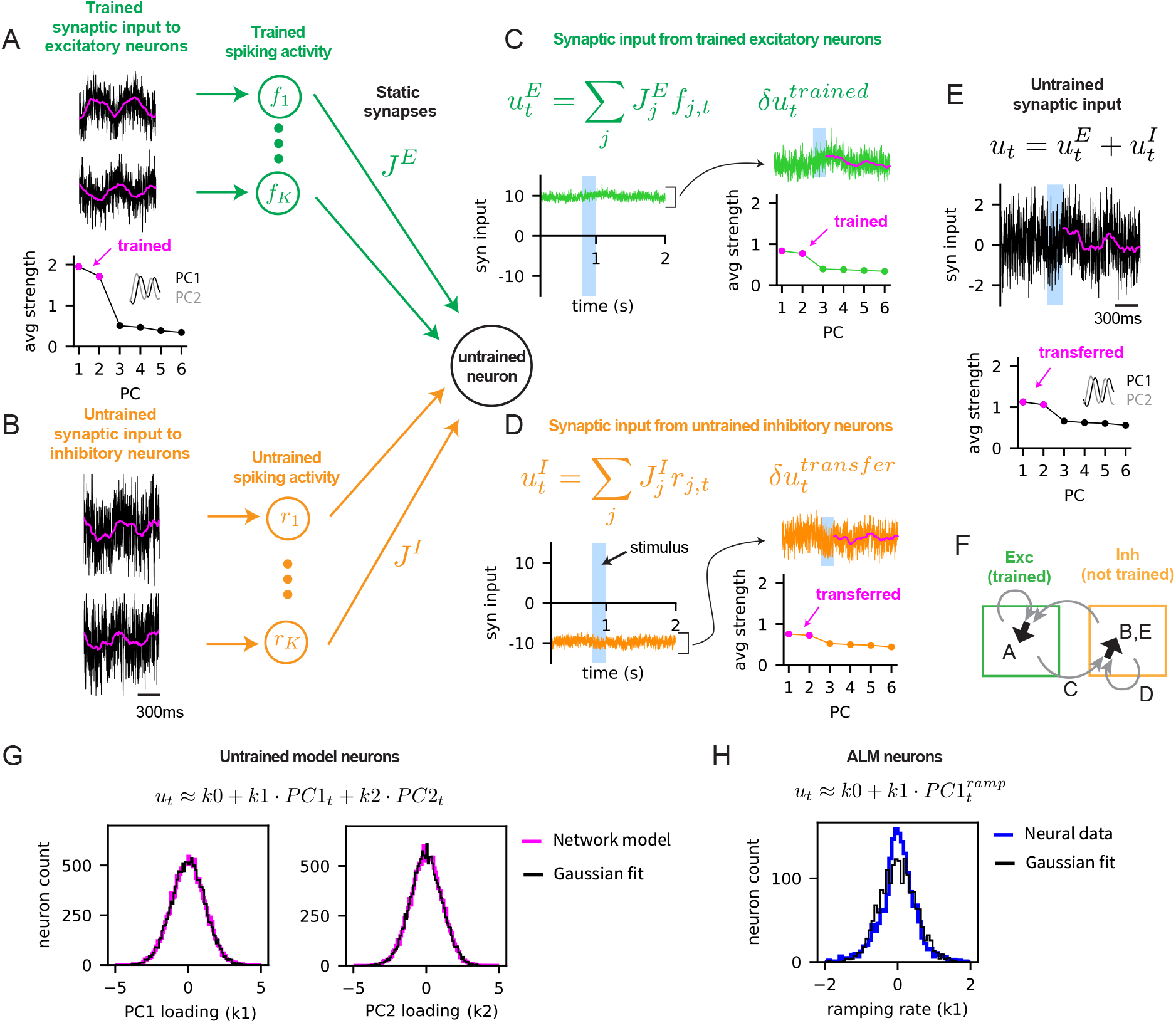
Network mechanism for distributing trained neural activity to untrained neurons through strong, non-specific connections. **(A)** Excitatory neurons were trained to generate 2Hz sinusoidal synaptic activity patterns with random phases. Examples (top) of trained synaptic inputs (black) to the excitatory neurons and their moving averages over 200ms window (magenta). The absolute value of the loading of each PC on trained synaptic activities (bottom) was averaged over all the excitatory neurons to obtain the average strength of the PCs. The first two PCs, which are the Fourier modes of 2Hz sine waves, are highlighted (magenta) and shown in the inset (PC1, PC2). *f_i_*’s in the circles (right) represent the spiking activity of trained neurons. Arrows (green) to an untrained neuron represent random, static, excitatory synapses with the synaptic weight *J_E_*. **(B)** Inhibitory neurons in the network were not trained. Examples of untrained synaptic inputs to inhibitory neurons (left). *r_i_*’s in the circles (right) represent the spiking activity of untrained neurons. Arrows (orange) to an untrained neuron represent random, static, inhibitory synapses with the synaptic weight *J_I_*. **(C)** Aggregate synaptic input from trained excitatory neurons to an untrained inhibitory neuron 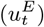 and its temporal modulation 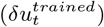 around the mean activity. The PCs (right) show the average strength of each PC in 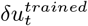. The PCs corresponding to the trained activity in panel (A) are highlighted (magenta). **(D)** Same as in (C) but for the aggregate synaptic input from untrained inhibitory neurons in the network to the same untrained inhibitory neuron shown in (C). **(E)** The total synaptic input (*u_t_* or the sum of 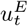 and 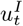) to the untrained inhibitory neuron (black) with the moving average (magenta). More examples are shown in panel (B). The PCs (bottom) show the strength of each PC in *u_t_*, averaged over all the untrained inhibitory neurons. The PCs corresponding to the trained activity in panel (A) are highlighted (magenta) and shown in the inset (PC1, PC2). **(F)** Schematic of synaptic inputs shown in panels (A) to (E). Total synaptic input to trained excitatory neurons (A: black arrow) is the sum of inputs from excitatory and inhibitory neurons (gray arrows). Total synaptic input to untrained inhibitory neurons (B,E: black arrow) is the sum of inputs from excitatory (C: gray arrow) and inhibitory neurons (D: gray arrow). **(G)** Distributions (magenta) of PC1 (*k*1) and PC2 (*k*2) loadings on the total synaptic input to the untrained inhibitory neurons (i.e., *u_t_* in panels (B) and (E)), overlaid with the Gaussian fits (black). The PCs are shown in panel (E), bottom. **(H)** Distribution (blue) of PC1 (*k*1) loadings on the estimated synaptic inputs to ALM pyramidal neurons for the lick-right trial type, overlaid with the Gaussian fit (black). The transfer function of the model neuron was used to estimate the synaptic input that yielded ALM neuron’s firing rate (see Methods). 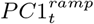 was a ramping mode, similarly to PC1 in Fig. 2E.

To explain the network mechanism underlying the spreading of trained activity patterns to the untrained neurons in the balanced regime, we considered a training setup where all the excitatory neurons were trained, while the inhibitory neurons were not. We chose the target activity to be 2Hz sine functions with random phases. After training, the temporal modulation of synaptic inputs to the excitatory neurons followed the target activity patterns (**Fig. 5A, top**). As a result, the first two PCs of the trained activities were 2Hz sine and cosine functions and were the dominant PCs of the trained activities (**Fig. 5A, bottom**).

The synaptic connections to an untrained neuron consisted of only the static synapses from randomly selected trained and untrained presynaptic neurons. Due to the large number of static synapses and their strong weights, the mean excitatory (**Fig. 5C**, 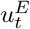) and inhibitory (**Fig. 5D**, 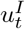) inputs to the untrained neuron were much larger, in absolute value, than their temporal modulations around the mean inputs (**Fig. 5C**, 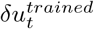; **Fig. 5D**, 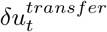). However, in the balanced regime, the large mean excitatory and inhibitory inputs dynamically canceled each other, resulting in the net mean input to the untrained neuron being around the spike-threshold (**Fig. 5E**, *u_t_*). Then, the spiking pattern of the untrained neuron was determined by the temporal modulation of the synaptic input around the net mean input, i.e., the sum of 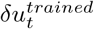 and 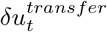.

To better understand how the trained activity patterns spread, we further examined the excitatory and inhibitory components of the temporal modulation of the synaptic input. Analysis of the input from trained excitatory neurons (**Fig. 5C**, 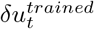) showed that its temporal modulation was dominated by the same PCs the excitatory neurons were trained to generate (**Fig. 5A**). The modulated input from trained excitatory neurons then led the total input to the untrained neuron to be modulated as well (**Fig. 5E**, *u_t_*). As a result, the untrained neurons produced modulated activity (**Fig. 5B**), which provided modulated input to neurons in the network (**Fig. 5D**, 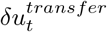). The temporal modulations of the total input to the untrained neurons (**Fig. 5E, bottom**) and the inputs the untrained neurons provided to other neurons (**Fig. 5D, right**) were both dominated by the same PCs acquired from training.

One of the predictions of this network mechanism is that the loadings of the PCs, at the level of the synaptic inputs to untrained neurons, are normally distributed. This is because the summation of a large number of randomly distributed PC loadings, determining the trained activity patterns, results in a Gaussian distribution when the synaptic weights are strong (see Methods). Indeed, this was the case for the loadings of the first two PCs in the example network trained on sine functions (**Fig. 5G**). Then we analyzed the loadings of the dominant PC mode in the ALM data, which were the slopes of the ramping activity of the synaptic inputs. Since the synaptic inputs to ALM neurons were not available, we estimated them by finding inputs to the transfer function of the model neuron that yielded the observed firing rates of ALM neurons. We found that the statistics of these loadings were also well fitted by a Gaussian distribution, supporting that the proposed mechanism may be biologically plausible (**Fig. 5H**).

The same network mechanism also provides an explanation for how functional properties, such as choice selectivity, can spread from neurons trained to be choice-selective to other neurons that are not trained (**Fig. 3E**). It stems from the fact that the differences in the activity of the lick-left and lick-right trials in the trained neurons spread through the random connections and are realized into two different responses in the untrained neurons, producing choice selectivity in them (see Methods for details). In addition, our mathematical analysis of the network mechanism is consistent with the findings that, due to their strong temporal modulations, inhibitory activity patterns spread more effectively than the excitatory activity patterns (**Fig. 4A**), and lower PC modes transfer with better fidelity than the higher PC modes (**Fig. 4D**; see Methods).

The results of our analysis show that the spreading of trained activity to untrained neurons is a general and robust circuit mechanism for strongly-coupled networks operating in the balanced regime. It is independent of the number of presynaptic inputs per neuron. Moreover, the slopes of ramping activity in the ALM neurons displayed statistics that agreed with the model prediction, providing an evidence for the biological plausibility of the proposed mechanism.

### Perturbation responses suggest that the ALM network operates in the balanced regime

We showed that when a subset of neurons was trained to reproduce the ALM activity, the task-related activity spread to the untrained neurons, which then also generated spiking activity resembling the ALM data (**Figs. 2, 3**). Such spreading of activity from trained to untrained neurons is a general mechanism for spreading activity in strongly coupled spiking networks (**Figs. 4, 5**). These results suggested that the observed task-related activity was produced in the ALM while it operated in the balanced regime. Here we used optogenetic perturbations to test if ALM activity displayed the characteristics of strongly coupled networks.

Specifically, we considered the activity modes of population of neurons responding to perturbations applied during the delay period (**Fig. 6A, B**). In strongly coupled networks consisting of excitatory and inhibitory populations, the projection of the population activity on the homogeneous mode (i.e. the average firing rate of the excitatory or inhibitory populations, **Fig. 6A**) is expected to recover rather fast from any perturbation. This is because the network dynamics of strongly coupled networks are extremely stable along the homogeneous mode [23]. To understand this phenomenon, one should consider changes to the average firing rate of the excitatory population in the network. This will result in a strong change (on the order of square root the number of inputs per neuron) to the excitatory drive to each of the neurons, which unless immediately suppressed by a strong inhibitory current, will destabilize the network. Therefore, to maintain the stability of network dynamics in strongly-coupled networks, a perturbation to the homogeneous mode is expected to decay quickly to its pre-perturbed value due to the strong and fast inhibition (a phenomenon known as ‘fast tracking’ [23, 44]).

**Figure 6:**
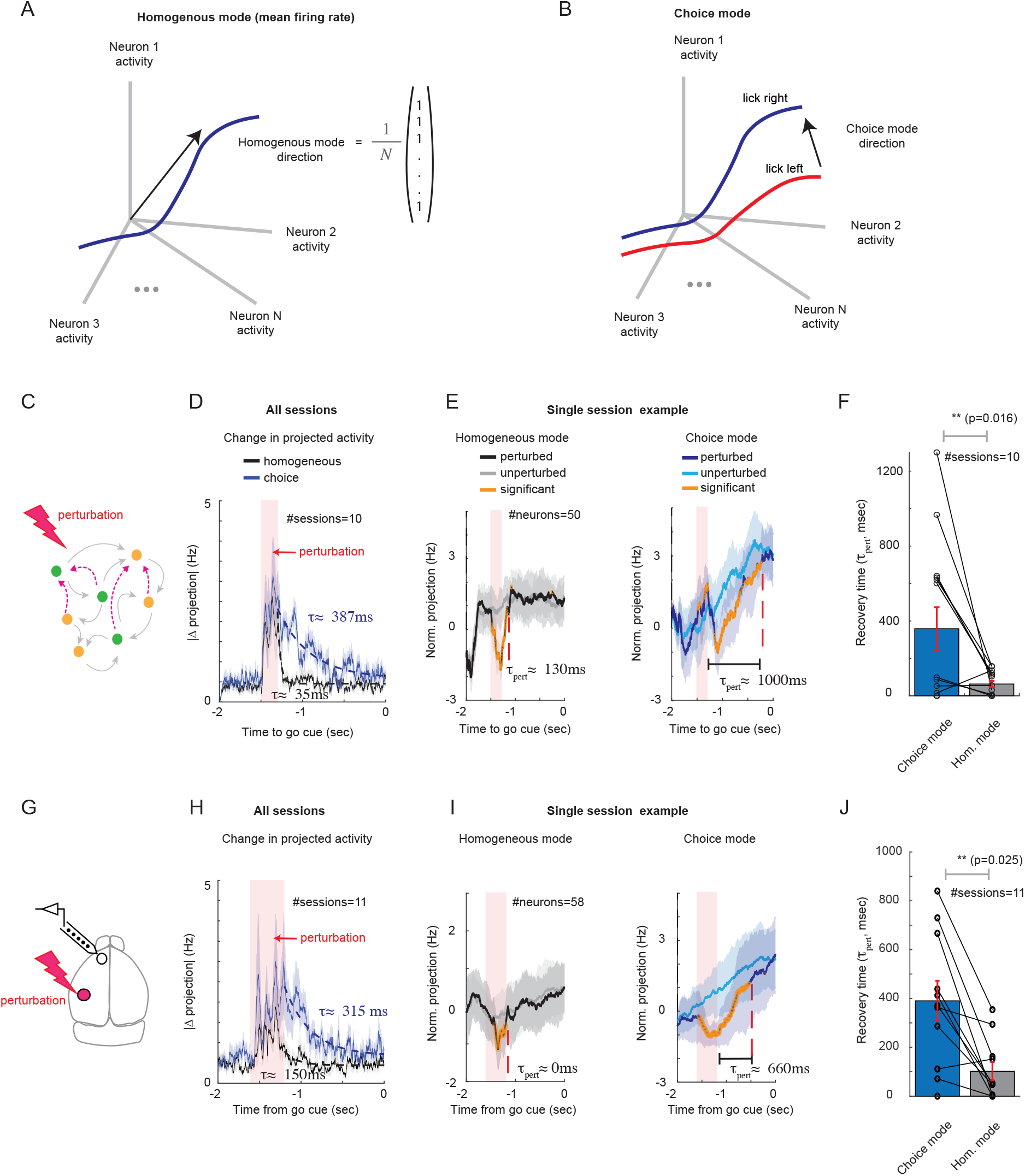
Fast and slow responses of the network to perturbations (model and data). **(A)** Schematic of the homogeneous mode, which averages the activity of the neurons. **(B)** Schematic of trial-averaged activity for lick-left (red) and lick-right (blue) trial types together with the choice mode in the neural activity space. This mode maximally separates trial-averaged activity with respect to licking directions (See Methods). **(C)** Schematic of a trained network receiving perturbation. **(D)** Change in projection on choice mode (blue) and homogeneous mode (black) against time, averaged over all 10 sessions. Each session consisted of sampling 50 neurons from the network. The change in projection was calculated as the trial-averaged activity for perturbed trials minus unperturbed trials (see Methods), with a 50ms smoothing. Mean± s.e.m. Shaded red: time of applied perturbation for perturbed trials. Dashed lines: exponential fit. **(E)** Projection of neural activity on homogeneous (left) and choice (right) modes for an example session, normalized by subtracting the average projection over the first 0.5 second of the delay period. Orange: significant differences between perturbed and unperturbed trials, starting from the perturbation time (see Methods). Dashed red: recovery time of perturbation, estimated as the first time the change was not significant following the perturbation (see Methods). **(F)** Recovery time for all sessions. Recovery of the homogeneous mode was significantly faster (p-value, by paired Student t-test) **(G)** Schematic of optogenetic perturbation in the mouse cortex. **(H)-(J)** Same as (D-F), but for putative excitatory neurons in ALM. Here each session corresponds to simultaneous recordings of ALM neurons on different days. Optogenetic perturbation in the data was applied to somatosensory cortex [21], whereas in the network model the stimulus that triggered the lick-left response was used to perturb the lick-right trials.

We verified this prediction by analyzing the responses of the motor cortex to optogenetic perturbations, and compared them with the responses of the trained strongly coupled network to similar perturbations. Consistent with our prediction, following a perturbation to the activity of neurons in the strongly coupled network (**Fig.2**), the projection on the homogeneous mode quickly returned to the baseline (**Fig. 6D, black**). In contrast, the projection of the activity on the choice mode (**Fig. 6B**), a mode that maximally separates trial-averaged activity with respect to licking directions (see [21, 41] and Methods), returned to the baseline after the perturbation with a significantly longer recovery time (**Fig. 6D, blue**; **Fig. 6F**, paired Student t-test, p-value=0.016). The slow recovery suggested that a dynamic attractor, which formed around the target trajectory due to training, was able to retract the perturbed activity at a slow timescale along the coding mode [21, 45]. Importantly, the network was trained only on the unperturbed ALM activity. Therefore, the fast and slow responses to perturbations were not dynamical properties acquired directly from the perturbed ALM activity, but instead they emerged from the strongly coupled network, when it was trained just on the unperturbed ALM activity.

To test the model prediction and verify that the ALM network operated in a similar dynamical regime, we conducted the same analysis on single sessions of simultaneously recorded ALM neurons (**Fig.6G-J**). We found that the response time of the homogeneous mode in ALM was significantly faster than that of the choice mode (**Fig. 6H-J**, paired Student t-test, p-value=0.025). Thus, the fast recovery of the homogeneous mode of ALM network, relative to the slow recovery of the choice mode, to optogenetic perturbations suggests that the ALM network operated in the same dynamical regime as the strongly coupled network. These findings suggest that the ALM network has the potential to be endowed with a network level mechanisms for generating widespread task-related activity, with limited synaptic reorganization on only a subset of neurons during learning.

## Discussion

In this study, we presented a circuit mechanism for distributing task-related activity in cortical networks. We have shown that neural activity learned by a subset of neurons can spread to the untrained parts of the network through pre-existing random connectivity, without additional training. This spread of activity occurs as long as the pre-existing random connections are strong and the network operates in the balanced regime. When a subset of neurons in the spiking network was trained to reproduce the activity of ALM neurons, the activity of untrained neurons in the network also displayed surprising similarity to the activity patterns of neurons in ALM. Single neuron activity patterns of the untrained neurons were ramping and selective to future choices, as was observed in ALM. Our work suggests that only a subset of neurons may be actively engaged in learning and the rest of the neurons are driven by the structured activity generated from the trained neurons.

Accumulating evidence show that inhibition in cortex is highly plastic (e.g. see review by [46]). We found that the fidelity of spreading the activity is higher when the inhibitory neurons are trained instead of the excitatory ones. For example, all of the excitatory neurons needs to be trained to explain 70% of the variance in the untrained inhibitory neurons, while training only 60% of the inhibitory neurons is enough to induce the same 70% variance in the untrained excitatory neurons (**Fig. 4A**). We speculate that this is a characteristic of the operating regime of cortical networks, in which typically the baseline spiking rates of inhibitory neurons is higher than the excitatory neurons. Inhibitory neurons can thus support stronger rate modulations (**Fig. 4B**), which in turn improves the fidelity of the spread (**Fig. 4A**, **Fig. 5**, Methods). Our results suggest that synaptic plasticity in inhibitory neurons can lead to wider spread of task-related activity in the motor cortex. Interestingly, this result is consistent with recent theoretical and computational studies showing that patterns of neural activity are primarily determined by inhibitory connectivity [47, 48].

In recent studies, the authors of [43, 49] argued that specific connectivity between excitatory and inhibitory neurons is necessary for choice selectivity to emerge in these two populations, based on computational models of their data. Our work suggests an alternative mechanism in which choice selectivity emerges in one population during training and spreads to the other population, without any reorganization of specific connections from the trained to the untrained populations. The network mechanism that spreads the task-related activity through random connectivity, as in our trained networks, is based on the susceptibility of neurons to modulations of synaptic inputs in strongly coupled networks (**Fig. 5**). This is a similar mechanism that explains how, without training or functional structure, orientation-selective neurons can emerge in primary visual cortex with a ‘salt-and-pepper’ organization [50, 51, 52].

While we studied the spreading of activity in recurrent spiking networks, a similar phenomena might generalize to other network architectures. For example, recent studies suggested that information can propagate through random connections in deep feedforward artificial networks [53, 54]. This suggests that the spreading of structured activity through unstructured connectivity is not restricted to local recurrent networks, but is applicable to other neural architectures and may provide the substrate for spreading information between brain areas.

Our study is related to and complements previous studies that trained rate-based units to reproduce neural activity in recurrent networks [19, 20, 21, 22]. In contrast to our spiking networks, each unit in these rate-based networks was paired with a target neuron activity for training, such that they could not investigate the effects of trained neurons on untrained neurons as we did here. We note that [19] (Supp. Fig. 3) examined what happened to the trained neurons when untrained neurons were included, but did not analyze the activity of untrained neurons. Furthermore, our network model consisted of spiking EI neurons operating in the balanced regime, and thus exhibited large temporal irregularity of spikes and trial-to-trial fluctuations, which is a key characteristic of cortical neurons. These properties stayed intact following learning. This contrasts with the commonly used rate-based chaotic networks that do not obey Dale’s law [55] and that suppress the trial-to-trial fluctuations that emerge due to the strong recurrent connectivity after training [35].

While our training algorithm requires only an order of square root of the pre-existing static connections to be plastic, the specific number of plastic connections may vary with the complexity of the trained neural activity. Indeed, almost three times more plastic synapses were needed to train the neurons to reproduce ALM activity, in which six PCs explained about 80% of the variance, compared to the number of plastic synapses needed to train sine waves, in which two PCs explained the same amount of the variance (**Table 2**). It is beyond the scope of this paper to determine exactly how the prefactor of the square root term depends on the complexity of the neural activity. However, previous studies suggest that the number of plastic connections might depend on the dimensionality (i.e., decay rate of the singular values) [56] or decorrelation time of the trained neural activity [37].

Several recent studies considered training spiking networks with dynamically balanced excitation and inhibition. In [57] the authors had to break the EI balance in order to achieve non-linear computations. With our training procedure, individual neurons can be trained to perform complex tasks, such as generating the spiking activity of cortical neurons, without leaving the balanced regime. The work by [58, 59] trained all the recurrent weights of the dynamically balanced spiking networks. To maintain strong excitatory-inhibitory activities after training, they considered weight regularizations that constrained the trained weights close to strong initial EI weights. Instead, in our training setup, the strong initial EI connections were left unchanged throughout training, thus always provided the strong excitation and inhibition. The trained neurons additionally received sparse plastic synapses, that were sparser than the initial EI connectivity. Such training scheme, which modifies only sparse synapses to trained neurons, is consistent with recent findings that build functional neural networks through weak, sparse, or low-dimensional synaptic modifications [19, 35, 60, 61, 62, 63]. It is also consistent with recent experimental evidence that hints for sparse, but functionally biased, synaptic connectivity in cortex [34].

There is an ongoing debate if cortex operates in the balanced regime [64]. Experimental evidence that are inconsistent with the balanced regime hypothesis mainly relies on data from sensory cortices. Here we present evidence that the motor cortex operates in the balanced regime by analyzing the recovery of neurons in the motor cortex to optogenetic perturbations. The presence of two recovery time scales, i.e., fast for the homogeneous mode and slow for the choice mode, is consistent with the prediction of the balance state on the ability of the homogeneous mode to rapidly track inputs, a phenomenon termed ‘fast tracking’ [23, 44]. Our analysis is different from the paradoxical effect observed in excitatory-inhibitory networks, where strong recurrent excitation must be compensated by strong feedback inhibition to maintain a stable network state [52, 65, 66, 67, 68].

We note that, with the use of moderate plastic inputs in our trained network, the plastic input was on the order of the spike-threshold. The network can thus implement non-linear computations at the individual neuron level. It can also support non-linear computations at the population level, as long as the computation is held by subpopulations, such that the overall excitatory and inhibitory population rates are unchanged after training [58, 62, 69]. Thus, the only mode that is strictly linear with the inputs to the network is the homogeneous mode. This is different from recent works that portrayed that the strict linear input-output relationship of balanced networks limits their computational power [57, 64].

To conclude, our work shows that while large changes in network dynamics can be observed during learning, attributing such changes to synaptic reorganization between neurons must be taken with care. In networks that operate in the balanced regime, in which motor cortex might operate, widespread changes in neuronal activity can be mainly a result of distributing learned activity from a dedicated subset of neurons to the rest of the network through strong but mostly unstructured connectivity.

## Acknowledgements

We would like to thank Larry Abbott and Sandro Romani for their valuable feedback. A.F., K.S. and R.D. were supported by the Howard Hughes Medical Institute. C.M.K. and C.C.C were supported by the Intramural Research Program at the NIDDK/NIH. C.M.K. would like to thank the Visiting Scientist Program at Janelia Research Campus for their support.

## Methods

### Data analysis

#### Principal component analysis of population rate dynamics

To obtain the PSTHs of neurons in a trained network, we repeated the simulation of a trained network 400 times starting at random initial conditions and applied the same external stimulus to trigger the trained activity patterns. Subsequently, for each neuron, the spikes emitted over multiple trials were placed in 20ms time bins, which ranged over the *T_target_* long training window, and averaged across trials to compute the instantaneous spike rates.

Given the PSTHs, 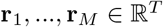, of a population of *M* neurons, we subtracted the mean rate of every neuron from its PSTH to remove differences in the baseline firing rates. In the following, we use the same notation **r**_*i*_ to refer to the mean subtracted PSTH of neuron *i*.

We then performed principal component analysis on the population rate dynamics **R** = (**r**_1_,…, **r**_*M*_), which is an *T* × *M* matrix, to obtain the principal components 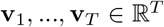 and the principal values *λ*_1_,…, *λ_T_*. This is equivalent to finding the eigenvectors and eigenvalues of the covariance matrix, **RR**^T^. The variance explained by the *k^th^* principal component was 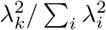.

The same procedure was applied to all the principal component analyses performed in this study.

#### Shared variance analysis

We identified population vectors along which the population activities of inhibitory model neurons and fast-spiking ALM neurons co-varied maximally [40]. We also quantified the fraction of variance that can be explained by the projected population-averaged activities (**Fig. 3**).

We first computed the correlation *C_ij_* = *corr*(**f**_*i*_, **g**_*j*_), which an *M*_1_ × *M*_2_ matrix, between the PSTH’s of inhibitory model neurons 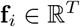, 1 ≤ *i* ≤ *M*_1_ and fast-spiking ALM neurons 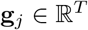, 1 ≤ *j* ≤ *M*_2_ where *M*_1_ = 2500, *M*_2_ = 306 and *T* = 100. Then the singular-value decomposition **C** = **UΣV** of the correlation matrix was performed, where **U** is an *M*_1_ × *M*_1_ matrix and **V** is an *M*_2_ × *M*_2_ matrix, to obtain the left singular vectors **U** = (**u**_1_,…, **u**_*M*_1__) with 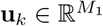 and the right singular vectors **V** = (**v**_1_,…, **v**_*M*_2__) with 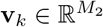.

To obtain the population-averaged activity along the singular vectors, the matrices of population rate, i.e., 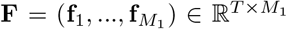 for the inhibitory model neurons and 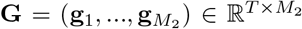 for the fast-spiking ALM neurons, were projected to the corresponding *k^th^* singular vectors **u**_*k*_ and **v**_*k*_, respectively, to obtain the *k^th^* shared components, 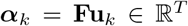 and 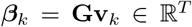. The variance explained by the *k^th^* shared component was defined as 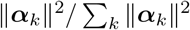 and 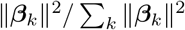, respectively.

#### Defining the choice and homogeneous modes

Trial-averaged spike rate of a neuron *i, r_i_*(*t,k*), were calculated for each trial, *k*, using 1ms bin size and were filtered with a 200ms boxcar filter.

We then analyzed the population dynamics of *N* simultaneously recorded neurons in a session. During each trial, the population activity of these neurons, **r**(*t, k*), drew a trajectory in the *N*-dimensional activity space. We identified the choice mode as *N* × 1 vector of trial-averaged spike rate differences of *N* neurons during trials with lick-right and lick-left outcomes, averaged within a 1sec window at the end of the delay epoch, before the go cue [21]:

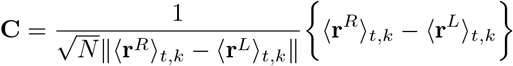

with the *L*_2_ norm, ║*x*║, and 〈*x*〉_*t,k*_ which is averaging over trials and time. The 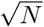 term was introduced to ensure that the projection of the neural activity is independent of the number of recorded neurons and for consistency with the homogeneous mode below. Projections of the neural activity along the choice mode were:

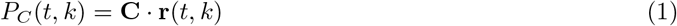

Similarly, the projection over the homogeneous mode was given by 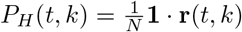, with **1** being a vector of ones.

If an individual neuron was not recorded during a particular trial, its weight in Eq.(1) was set to zero, and for the analysis we selected trials with at least 10 simultaneously recorded neurons.

#### Response of the modes to perturbations

To assess the impact of vS1 photostimulation during the delay on the homogeneous and choice modes in the ALM, we computed for each session the single-trial projections on each of the modes, *P_C_*(*t, k*) and *P_H_*(*t, k*), for correct lick-right trials both with and without the photostimulation. The trial-averaged activity was plotted for one example session in **Fig. 6E,I** along with the SD, after subtracting the average projection over the first 0.5 seconds of the delay period.

We used a statistical hypothesis test (Student t-test) to estimate the decay time back to the non-perturbed trajectories for the projections on the modes. Specifically, for each time bin we tested the null hypothesis that the perturbed and unperturbed trials were from the same distribution and rejected the null hypothesis with a p-value< 0.05 (orange dots in **Fig. 6E,I**). We only analyzed sessions in which the photostimulation resulted in a significant change in at least 10% of the time points during the photostimulation period ([-1.6,1.2]sec, 13/17 sessions). To calculate the decay time, we then used the last significant time bin within the time window of [-1.2, 0] sec for which the derivative was smaller than 10ms (dashed red lines in **Fig. 6E,I**). The perturbations in 2/13 sessions were biased and were not included in the analysis, leaving 11 sessions of simultaneously recorded neurons.

To calculate the decay time over all sessions (**Fig. 6D**) we averaged the projection in each of the 11 analyzed sessions and calculated the difference in the projection between the perturbed and unperturbed trials (Δ projection). We then took the absolute value and averaged over all sessions (**Fig. 6D**, mean± SEM). Finally, we estimated the decay rate by an exponential fit.

### Spiking neural networks

#### Network connectivity

The spiking neural network consisted of randomly connected *N_E_* excitatory and *N_I_* inhibitory neurons. The recurrent synapses consisted of static weights **J** that remained constant throughout training and plastic weights **W** that were modified by the training algorithm. The static synapses connected neuron *j* in population *β* to neuron *i* in population *α* with probability *p_αβ_* = *K_αβ_/N_β_* and synaptic weight 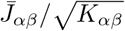, where *K_αβ_* is the average number of static connections from population *β* to *α*:

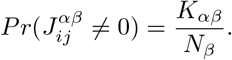

The strength of plastic synapses, 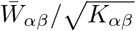, was of the same order as the static weights. However, the plastic synapses connected neurons with a smaller probability:

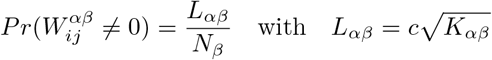

which made the plastic synapses much sparser than the static synapses [62]. Here, *c* is an order 1 parameter that depends on training setup.

The static and plastic connections were non-overlapping in that any two neurons in the network can have only one type of synapse.

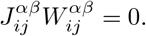

Keeping them disjoint allowed us to maintain the initial network dynamics generated by the static synapses and, subsequently, introduce trained activity to the initial dynamics by modifying the plastic synapses.

The static recurrent synapses were *strong* in that the coupling strength between two connected neurons scaled as 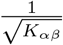, while the average number of synaptic inputs scaled as *K_αβ_*. This is in contrast to the weak, 1/*K_αβ_*, coupling we considered in **Fig. S9**. As a result of this strong scaling, the excitatory 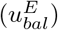 and inhibitory 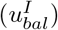 synaptic inputs to a neuron from static synapses increased as 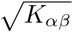, thus were much larger than the spike-threshold for a large *K_αβ_*. However, 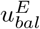 and 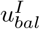 were dynamically canceled, and the sum (*u_bal_*) was balanced to be around the spike-threshold ([23], **Fig. 1B, middle**).

In contrast to the static synapses, each trained neuron received only about 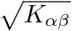 plastic synapses. This made the plastic synapses much sparser than the sparse static EI connectivity (e.g., with *K* = 1000 static synapses, there are of the order of 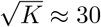 plastic synapses per neuron). Consequently, the EI plastic inputs 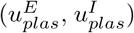 of the initial network were independent of *K_αβ_* and substantially weaker than the EI balanced inputs 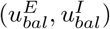 for a large *K_αβ_*. After training the plastic synapses, the total synaptic input (*u* = *u_bal_* + *u_plas_*) to each trained neuron was able to follow the target patterns (**Fig. 1B, left**; **Fig. 1C**), while the plastic input (*u_plas_*) stayed around the spike-threshold (**Fig. 1B, right**). With this scaling of plastic synapses, training was robust to variations in the number of synaptic connections, *K_αβ_*. Network trainings were successful even when *K_αβ_* was increased, such that the excitatory and inhibitory balanced inputs were tens of orders of magnitude larger than the plastic inputs (**Fig. S1**).

From a technical point of view, the choice to train only a very sparse number of plastic synapses made the plastic inputs to be on the order of the spike-threshold (i.e. order one, independent of the number of connections). Therefore, they could not affect the mean firing rates of the excitatory and inhibitory neurons in the network. These were fully determined by a linear equation, termed the ‘balanced equation’ [23, 70], that only involves the strength of the static connections, 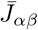, and the external inputs. Alternatively, if the plastic synapses were more abundant in the network, on the order of the number of static connections, they could interfere with the the ability of the strong inhibition to balance the strong excitation for each neuron in the network. Such interference significantly limits the ability to train the spiking networks. Taking the number of plastic connections to be on the order of 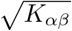 thus allowed to both train the networks, and keep it in the balanced regime.

#### Network dynamics

We used integrate-and-fire neuron to model the membrane potential dynamics of the *i*’th neuron:

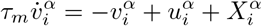

where a spike is emitted and the membrane potential is reset to *v_reset_* when the membrane potential crosses the spike-threshold *v_thr_*.

Here, 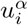 is the total synaptic input to neuron *i* in population *α* that can be divided into static and plastic inputs incoming through the static and plastic synapses, respectively:

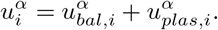

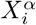 is the total external input that can be divided into constant external input, plastic external input, and the stimulus:

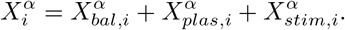

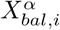 is a constant input associated with the initial balanced network. It scales with the number of connections, i.e., proportional to 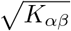, determines the firing rate of the initial network and stays unchanged [23]. 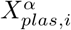 is plastic input provided to trained neurons in the recurrent network from external neurons that emit stochastic spikes with pre-determined rate patterns. The synaptic weights from the external neurons to the trained neurons were modified by the training algorithm. 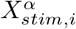 is the pre-determined stimulus, generated independently from the Ornstein-Ulenbeck process for each neuron, and injected to all neurons in the network to trigger the learned responses in the trained neurons.

The synaptic activity was modeled by instantaneous jump of the synaptic input due to presynaptic neuron’s spike, followed by exponential decay. Since the static and plastic synapses did not overlap, we separated the total synaptic input into static and plastic components as mentioned above:

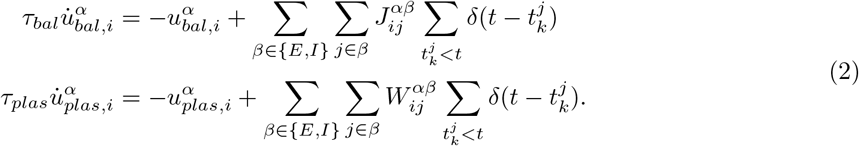

Alternatively, the synaptic activity can be expressed as a weighted sum of filtered spike trains because the synaptic variable equations (Eq. 2) are linear in **J** and **W**:

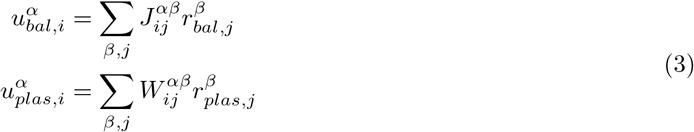

where

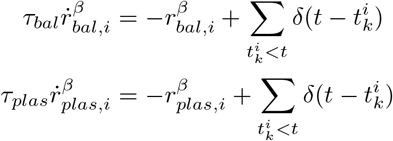

describe the dynamics of synaptically filtered spike trains.

Each external neuron emitted spikes stochastically at a pre-defined rate that changed over time. The rate patterns, followed by the external neurons, were randomly generated from an Ornstein-Ulenbeck process with mean rate of 5 Hz. The synaptically filtered external spikes were weighted by plastic synapses **W_X_** and injected to trained neurons:

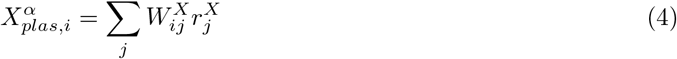

where

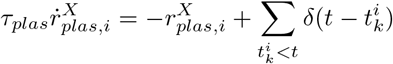

Similarly, the external stimulus *X_stim,i_* applied to each neuron i in the network to trigger the learned response is generated independently from the Ornstein-Ulenbeck process.

In the following section, we will use the linearity of **W**, **W_X_** in Eqs. 3 and 4 to derive the training algorithm that modifies plastic synaptic weights.

### Network training scheme

#### Overview

Prior to training the network, neurons were connected by the recurrent static synapses and emitted spikes asynchronously at constant rates. This asynchronous state of the initial network has been investigated extensively in previous studies [23, 25, 70].

Starting from this asynchronous state, the goal of training was to produce structured spiking rate patterns in a subset of neurons selected from the network. Specifically, our training scheme modified the recurrent and external plastic synapses projecting to the selected neurons, so that they generated target activity patterns when evoked by a brief external stimulus. To this end, we first selected *M* neurons to be trained from a network consisting of *N* neurons, and then prepared *M* target functions *f*_1_(*t*),…, *f_M_*(*t*) defined on a time interval *t* ∈ [0, *T_target_*] to be learned by the selected neurons. The plastic synapses projecting to each selected neuron *i* were then modified by the training algorithm such that the total synaptic input *u_i_*(*t*) to neuron *i* followed the target pattern *f_i_*(*t*) on the time interval *t* ∈ [0, *T_target_*] after the training.

#### Initialization of plastic synapses

For each trained neuron, we randomly selected *L* excitatory and *L* inhibitory presynaptic neurons that projected plastic synapses to the trained neuron. When the excitatory subpopulation was trained, the presynaptic excitatory neurons were sampled from other trained excitatory neurons while the presynaptic inhibitory neurons were sampled from the entire inhibitory population. Similarly, when the inhibitory subpopulation was trained, the presynaptic inhibitory neurons were sampled from other trained inhibitory neurons while the presynaptic excitatory neurons were sampled from the entire excitatory population. The untrained neurons did not receive any plastic synapses.

Each trained neuron also received inputs from all the *L_X_* external neurons. The plastic weights from the external neurons to each trained neuron were trained by the learning algorithm.

#### Cost function

Each trained neuron i had its own private cost function defined by

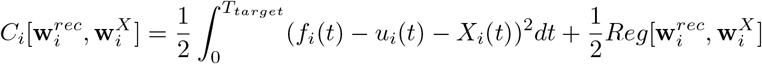

where 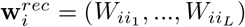 is a vector of recurrent plastic synapses to neuron *i* from other presynaptic neurons in the network indexed by *i*_1_,…, *i_L_*. Similarly, 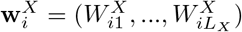 is vector of plastic synapses to neuron *i* from the external neurons. The regularization of plastic weights 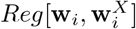 consisted of two terms

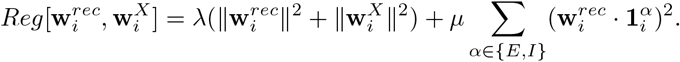

The first term is a ridge regression that evaluates the *L*_2_-norm of the plastic weights. The second term is called ROWSUM regularization where the elements of the vector 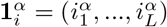 are defined to be 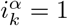 if the presynaptic neuron *i_k_* belongs to population *α* and 0 otherwise [59]. The inner products 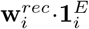 and 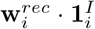 are the aggregate plastic weights to neuron i from the excitatory and inhibitory populations, respectively. Including the ROWSUM regularization allowed us to keep the aggregate excitatory and inhibitory plastic weights fixed throughout the training.

#### Training algorithm

We derived a synaptic update rule that modified the plastic synapses to learn the target activities. The learning rule was based on recursive least squares algorithm (RLS) that was previously applied to train the read-outs to perform tasks [35, 36] and the individual neurons to generate target activity patterns [19, 37, 45]. The derivation presented here closely follows previous papers [37, 59]. For notational simplicity, we dropped the index *i* in **w**_*i*_ and other variables, e.g., *f_i_, u_i_*. We note that the same synaptic update rule was applied to all the trained neurons.

The gradient of the cost function with respect to the vector of full plastic weights **w** = (**w**_*rec*_, **w**_*X*_) was

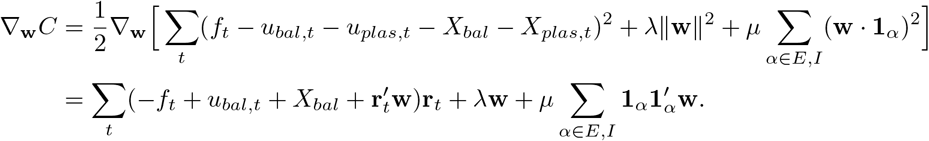

Here we substituted the expressions *u_plas,t_* = **w**_*rec*_ · **r**_*plas,t*_ and 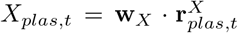 in the first line to evaluate the gradient with respect to **w**. In the second line, we used a condensed expression 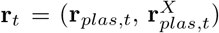 to denote the synaptically filtered spike trains from all plastic inputs. The vectors **1**_*α*_ apply only to the recurrent plastic weights **w**_*rec*_ and take zero elements on **w**_*X*_.

To derive the synaptic update rule, we computed the gradient at two consecutive time points

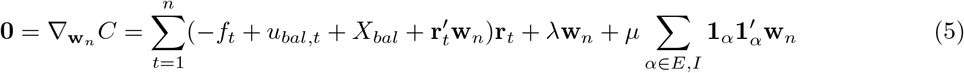

and

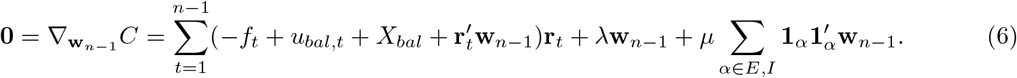

Subtracting Eqs (5) and (6) yielded

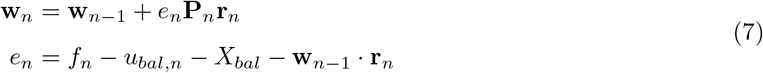

where

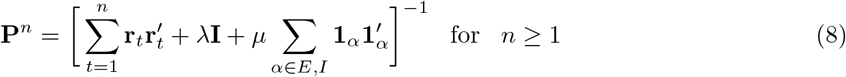

with the initial value

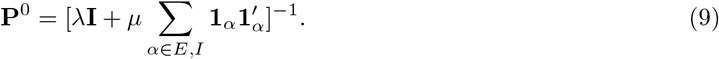

To update *P^n^* iteratively, we used the Woodbury matrix identity

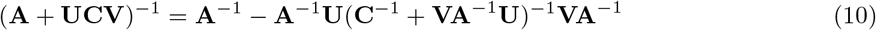

where **A** is invertible and *N* × *N*, **U** is *N* × *T*, **C** is invertible and *T* × *T* and **V** is *T* × *N* matrices. Then **P**^*n*^ can be calculated iteratively

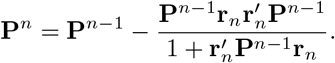

#### External stimulus triggering target activity patterns

To trigger the target activity patterns learned by the trained neurons, a brief external stimulus (200ms long) was applied to every neuron in the network immediately before generating the activity patterns. Two different sets of stimuli were prepared to trigger the lick-left and lick-right population responses. One set of stimulus was used during and after training to trigger the lick-left response and the other set of stimulus was used for the lick-right response. The stimulus 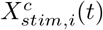 to each neuron *i* and trial type *c* = *L,R* was generated independently from the Ornstein-Ulenbeck process: 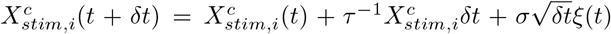 where *τ* = 20ms, *σ* = 0.2 and *ξ*(*t*) was uncorrelated Gaussian distribution with zero mean and unit variance.

#### Generating sinusoidal activity patterns

For demonstrating the Subset Training method (**Fig. 1**) and the network mechanism for spreading trained activity (**Fig. 5**), neurons in the network were trained to follow sine functions with random phases. Specifically, neuron *i* in the network learned the target pattern *f_i_*(*t*) = *a*sin(*ωt* + *ϕ_i_*) + *b_i_* on the time interval *t* = [0,1]sec, where the amplitude *a* = 0.5, the frequency *ω* = 1*rad/sec* (Fig. 1) and 2*rad/sec* (Fig. 5), the phase *ϕ_i_* was sampled from a uniform distribution [0, 2*π*], and the offset *b_i_* was the mean synaptic input to the neuron in the initial balanced network prior to training.

#### Generating target neural trajectories

A subset of excitatory neurons in the network learned to reproduce the PSTHs of pyramidal neurons recorded from ALM in [21]. For each pyramidal neuron, the spikes emitted across multiple experiment trials were placed in Δ*t* = 20ms time bins that ranged over the *T_target_* = 2 second delay period. The PSTHs were then smoothed by a moving average over a 300ms time window centered at each time bin. We obtained two sets of PSTHs 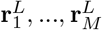 and 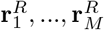 from *M* = 1824 pyramidal neurons for the lick-left and lick-right trial types. Each PSTH 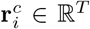 for neuron *i* and trial-type *c* ∈ *L,R* was an *T* = *T_target_*/Δ*t* = 100 dimensional vector defined on time points *t* = [−2 + Δ*t*,…, −Δ*t*, 0]sec, where 0 is the onset of go-cue.

Next, we converted the PSTHs to target synaptic activities to be used for training the synaptic inputs to selected neurons. For each spike rate 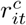 where *i* = 1,…, *M*, *c* = *L, R* and *t* = –2 + Δ*t*,…, 0, we obtained the mean synaptic input 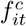 that needs to be applied to the the leaky integrate-and-fire neuron to generate the desired spike rate. To this end, we numerically solved the nonlinear rate equation

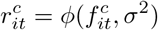

where 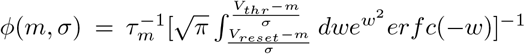 is the transfer function of the leaky integrate-and-fire neuron given mean input, *m*, and variance of the input, *σ*^2^ [27, 71]. We obtained the synaptic fluctuation *σ* from the synaptic noise in the neurons of the initial network since the slow plastic inputs did not significantly change the fast noise fluctuation. This conversion yielded two sets of target synaptic inputs 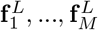 and 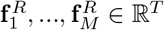 for *M* excitatory neurons to be trained.

We chose the parameters of the initial network connectivity such that the mean rate of the excitatory and inhibitory populations in the network was close to estimated mean rates of the ALM data (mean excitatory rate was 4.2 Hz and inhibitory rate was 11.0 Hz). To select the subset of excitatory neurons to be trained, we compared the mean firing rates of the neurons in the initial network with the firing rates of pyramidal neurons and identified the excitatory neuron whose firing rate’s was closest to the pyramidal neuron. This process was repeated until all the pyramidal neurons were matched to the excitatory neurons uniquely.

#### Generating target synthetic trajectories

To generate synthetic data that shared similar statistics and low-dimensional dynamics as the neural data, we performed PCA on the PSTHs of pyramidal ALM neurons to identify the principal components 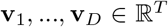 that explained majority of their variance. We found that *D* = 9 was large enough to explain over 95% of the variance. The same procedure was applied to the PSTHs of the fast-spiking ALM neurons to obtain their principal components.

We sought to construct synthetic trajectories 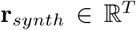 that resembled the PSTHs of the pyramidal and fast-spiking ALM neurons (**Fig. S6**). To this end, we expressed the synthetic trajectory **r**_*synth*_ as a weighted sum of the principal components: 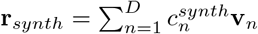. To find the distribution of the coefficients 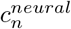 of the neural data, we projected the PSTHs of pyramidal neurons onto the PCs and obtained the empirical distribution of 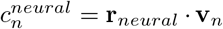. Bootstrapping the synthetic coefficients 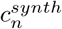 from the empirical distribution of 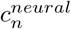 was performed in two steps. First, the mean firing rate of synthetic target was sampled from the empirical rate distribution to generate synthetic PSTHs that had rate distribution statistically identical to the empirical distribution (**Fig. S6A, B**). Next, since 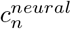 depended strongly on the mean firing rate of neurons, 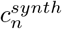 was bootstrapped from a subset of 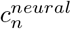 whose underlying firing rate was close to the firing rate of synthetic target (**Fig. S6C**). In this way, the distributions of the firing rates and PC loadings of the synthetic and neural data were almost identical (**Fig. S6E**).

In addition, we generated the synthetic PSTHs in pairs for the lick right and lick left trials. First, the PSTHs for the lick right and lick left conditions were generated independently. Then, we sorted the PSTH’s of each condition separately and paired them, to ensure the pairs had similar level of mean firing rates. Subsequently, we added Gaussian noise with zero mean and standard deviation equal to the difference of lick right and lick left mean firing rates, to the PSTH’s of the lick left condition. This allowed us to introduce choice selectivity to the synthetic PSTHs.

The synthetic PSTHs were then converted into target synaptic inputs following the same procedure applied to the neural PSTHs.

### Mathematical analysis of the effects of trained inputs on untrained neurons

In this part of the methods we use mathematical analysis to show how random inputs from trained neurons can drive the untrained neurons to follow the trained activity, without further training, if the network operates in the balanced regime.

To simplify the analysis, we assumed that only the excitatory population was trained and the inhibitory population was not. In addition, we assumed that the target functions, *f_it_* for neuron *i* and *t* ∈ [0, *T_target_*], were slower than the slow plasticity signal and that training was perfect. In this case, we can approximate the total synaptic input to a trained excitatory neuron using the fixed point equation:

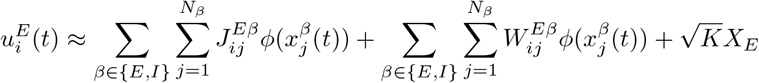

with 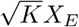 the strong external input associated with the balanced network [23]. The transfer function, 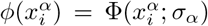, was the Riccardi function [27, 71], with 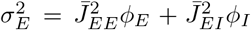. The population rate was given by 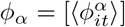, with 〈*x*〉 denoting the average over the time and [*x*] the average over the neurons.

Similarly, the total synaptic input to an untrained neuron, which lacked plastic connections, followed:

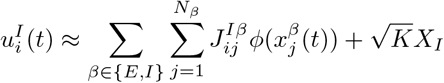

with 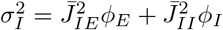.

Our goal was to analyze the synaptic drive from the trained (excitatory) neurons to untrained (inhibitory) neurons to make specific predictions about what aspects of the trained inputs allowed them to spread effectively to the untrained neurons.

#### Statistics of random inputs from the trained neurons to an untrained neuron

If an excitatory neuron i is successfully trained, its firing rate closely follows the target activity *f_it_*. We used a shorthand notation 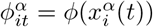 and expressed the firing rate of the trained neuron in the form 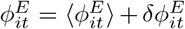, with the temporal modulation 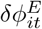. We next considered the singular value decomposition of the temporal modulation:

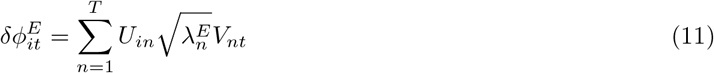

which is *N* × *T* matrix, and where **U** is a *N* × *N* matrix of the left singular vectors and **V** is *T* × *T* matrix of the right singular vectors. Here, we considered a discretized version of time with *T* = *T_target_*/Δ*t*, such that the matrices are of finite size. The values 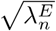 are the singular values (SVs) and 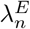 are the elements of the spectrum of the covariance matrix of the trained excitatory neurons. For instance, if we choose the target activity to be sinusoidal functions with random phases (**Fig. 5A**), the covariance matrix is stationary and the right singular vectors are the Fourier modes (e.g., *V*_1*t*_ ∝ sin(*ωt*), *V*_2*t*_ ∝ cos(*ωt*)).

Untrained (inhibitory) neurons do not receive plastic synapses. Thus, the aggregate input from the trained neurons to an untrained neuron, 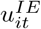, is a random summation of trained neurons’ activity. It is given by:

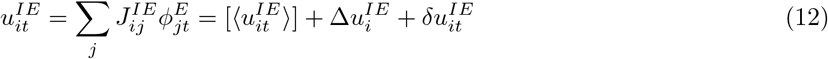

with the average population input 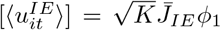. The second term in Eq.(12) is the quenched disorder [44] and its variance is given by:

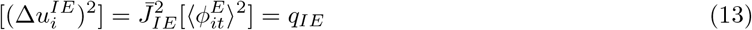

The last term in Eq.(12) is the temporal modulation of the aggregate trained input, 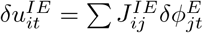. Using Eq.(11), it is given by:

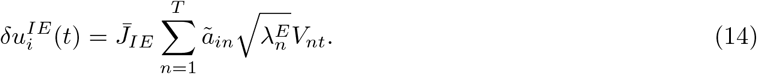

Here, due to the Central Limit Theorem, the coefficients 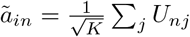 are Gaussian vectors with zero mean and unit variance in the large *K* limit (see **Prediction 1** below), if the left singular vectors *U_in_*’s are random variables with zero mean and unit variance. Importantly, we emphasize that it is the strong coupling (i.e., synaptic weights scale as 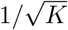) that allows the coefficients *ã_in_*’s to have finite variance. This is not the case if synaptic weights are weak (see **Prediction 2** below). In addition, the variance of the coefficients of temporal modulation is 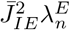, which shows that the SVs, 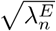, determine the strength of temporal modulation (see **Prediction 3** below).

With this, the synaptic input to an untrained neuron from the trained population can be written in the following form:

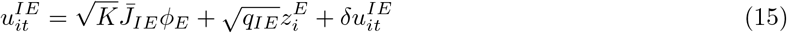

with 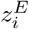 being a Gaussian random variable with zero mean and unit variance.

For example, when the target functions are sinusoidal functions with random phases (**Figs. 1, 5**) these temporal modulations are:

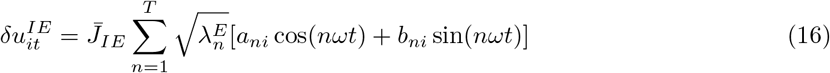

where we replaced *ã_ni_* in Eq.(14) with the even and odd coefficients of the cosine and sine functions, *a_ni_,b_ni_*, respectively.

Similarly, in the case of the ALM data, the dominant right singular vector is a ramping mode (**Fig. 2E,F**), i.e. *V*_1*t*_ ∝ *t* and the temporal modulations are dominated by:

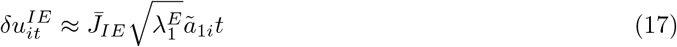

with 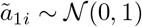.

#### The recurrent untrained inputs and implications

The synaptic input to an untrained inhibitory neuron consists of a large, 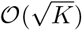, and positive mean drive from the excitatory neurons. The untrained neurons will thus fire with high rates and regular spiking, unless the network operates in the balanced regime, in which the recurrent inhibition cancels most of this large excitatory drive [23]. In this case, the untrained neuron will be driven by the temporal modulations originating from the random summation of the activity of trained neurons, which are of 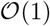 due to the strong coupling. This input is spanned by the principal components (or, equivalently, the right singular vectors) of the trained population according to Eq.(14).

A similar analysis on the recurrent inputs from the untrained inhibitory population, 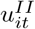, needs to be done to infer the statistics of the temporal fluctuations of the net input, 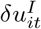, and rates, 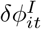, of the untrained inhibitory neurons. This analysis needs to be done in a self-consistent way to determine the statistics of 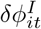 [56]. While this analysis is beyond the scope of the current paper, several observations can be made already by examining the statistics of the inputs from the trained population.

##### Prediction 1.

No matter what the right singular vectors (which we refer to as the PCs in the main text) are, their coefficients are expected to be Gaussian. This prediction is shown in **Fig. 5F** for artificial target functions of sine functions with random phases, as well as in **Fig. 5G** for the coefficients of the dominant ramping mode in the neural data.

##### Prediction 2.

The spread of activity in the network is possible only because the variance of *ã_it_*’s in Eq.(14) is finite. It is a result of the strong coupling in the network, i.e. the 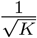 scaling of the synapses, which guarantees, due to the Central Limit Theorem, that the variance of the aggregate input from the trained neurons converge is finite. This is in contrast to the case of weak synapses (e.g., scaling of 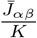 instead of 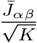), where the variance of *ã_it_* converges to zero in the large *K* limit (**Fig. S9**, no spreading of trained activity in a weakly coupled network).

##### Prediction 3.

The strength of the transfer of the trained activity to the untrained neurons depends on the variance of the trained population through Eq.(14). As shown in **Fig. 4B**, in the ALM data the variance of the temporal modulations of the inhibitory neurons is larger than those of the excitatory neurons. This result suggests why the fidelity of the spread improved when the inhibitory population was trained instead of the excitatory population. It also explains why lower mode PCs of the activity can spread better in the network, as their corresponding SVs (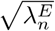 in Eq.(14)) are, by definition, larger than those of the higher mode PCs.

##### Prediction 4.

This framework provides additional insights into how excitatory neurons trained to be choice-selective can impart the learned selectivity to the untrained inhibitory neurons through nonspecific, strong synaptic connections (see **Fig. 3E**). To show this, one can estimate the statistics of the difference in the input to an untrained inhibitory neuron from the trained population for the lick-right and lick-left trials. For instance, if we consider the target functions to be defined by the dominant ramping mode that captures over 70% of the variance (**Fig. 2E**), the relevant basis function would be *V*_1*t*_ ∝ *t* for *t* ∈ [0, *T_target_*], and the selectivity of the trained inputs (*SI^E^*) yields

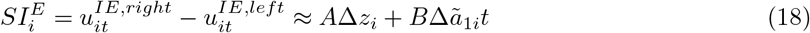

where *A* and *B* determine the variance in the baseline inputs and ramping rates, respectively. From Eq.(15), the quenched disorder yields Gaussian variables 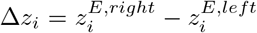, with a finite variance *A*^2^. From Eq.(17), the temporal modulation yields a Gaussian variables 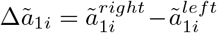, with a finite variance *B*^2^. Because Δ*z_i_* and Δ*ã*_1*i*_ are random variables with finite variances, the trained inputs develop choice selectivity, which can then elicit choice selectivity in the untrained inhibitory neurons (**Fig. 3D**). The good agreement of the distribution of choice-selectivity in the untrained neurons in the model and the putative fast-spiking neurons in the neural data (**Fig. 3D**) is consistent with this prediction.

### Simulation parameters

## Supplementary material

**Figure S1:**
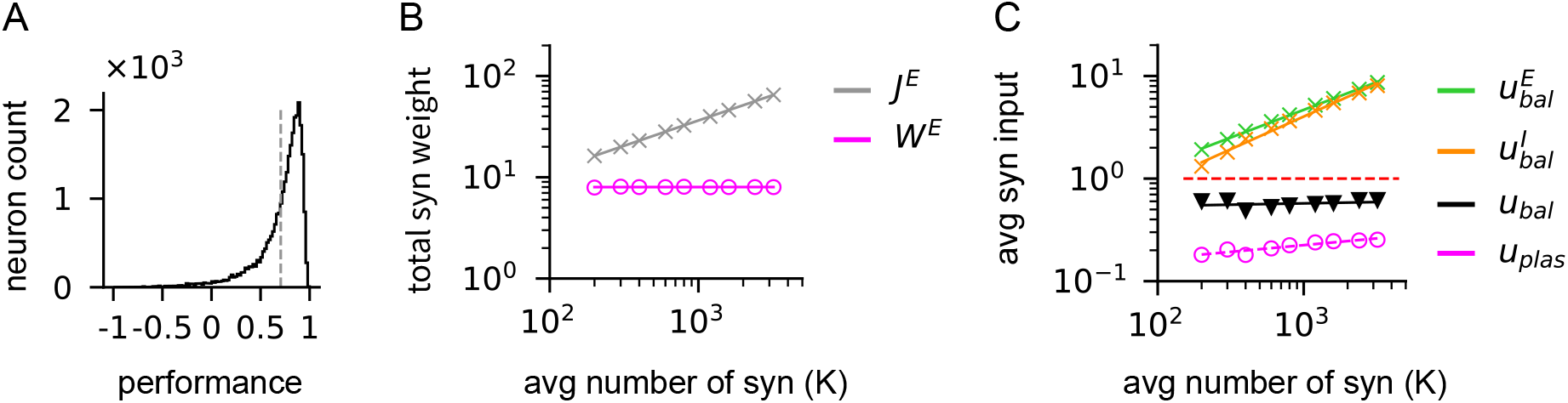
Training is robust to scaling of synaptic weights and inputs. All neurons in a network consisting of *N* = 30, 000 excitatory and inhibitory neurons were trained to learn sine functions with random phases. The average number of static synapses to a neuron, *K*, in the initial network was varied over a wide range to demonstrate that the training scheme is robust to the increased strength of static synapses. **(A)** Typical performance of neurons in a trained network. Average performance in dashed line. Here, the performance was quantified as the correlation between the target patterns and the synaptic input to a neuron that learned the target. **(B)** The number of plastic synapses was increased proportionally to 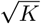 and the strength of plastic synapse by 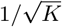. Therefore, the total weight of plastic synapses to a neuron remained constant in the initial network, independent of *K*(*W^E^*: total weight of plastic excitatory synapses to neurons). In contrast, the total weight of static synapses to a neuron increased as 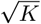 ([44]; *J^E^*: average weight of static excitatory synapses to neurons). **(C)** Strength of different types of synaptic inputs to neurons in networks trained over a wide range of *K*. For each *K*, networks were trained until the network performance reached 0.6. Trained networks showed that learning was successful even as the excitatory 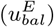 and inhibitory 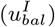 balanced inputs coming through the static synapses increased with 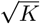, while the plastic inputs (*u_plas_*) coming through the plastic synapses and total balanced inputs 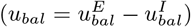 remained constant, on the order of threshold (dashed red line), independent of *K*.

**Figure S2:**
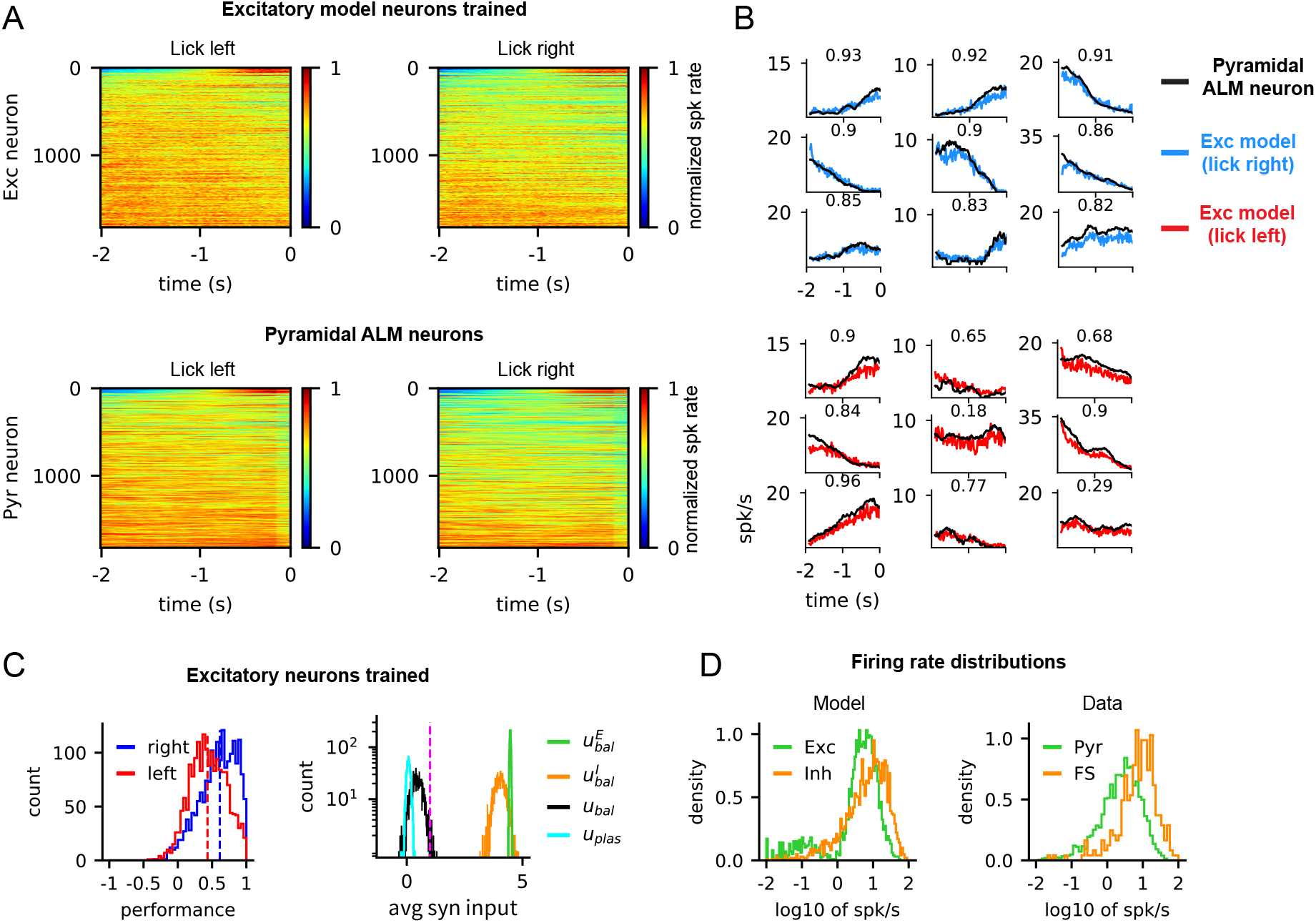
Spiking activity of excitatory neurons trained to reproduce the PSTHs of pyramidal ALM neurons. **(A)** Normalized PSTHs of trained excitatory neurons (top) and the normalized PSTHs of pyramidal ALM neurons (bottom) that were used to train the excitatory neurons. Neurons were sorted in decreasing order according to their performance in the lick right trials. **(B)** PSTHs of trained excitatory and pyramidal ALM neurons were compared for the lick-right (top) and lick-left (bottom) trials. Panels in the same position represent the activity of same neurons during the lick-right and lick-left trials. The correlation between two PSTHs are shown on each panel. **(C)** Performance (left) of all the trained excitatory neurons (average performance in dashed lines), quantified as the correlation between the PSTHs of the trained neuron and target ALM neuron. Distribution of mean synaptic inputs (right) to trained excitatory neurons. Here, 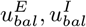 are the excitatory and inhibitory balanced inputs, respectively, to the trained neurons through the static synapses. *u_bal_* is the sum of 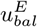 and 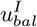. *u_plas_* is the plastic input to the trained neurons through the plastic synapses. **(D)** Distribution of the firing rates of neurons in a trained network (left) and ALM neurons (right).

**Figure S3:**
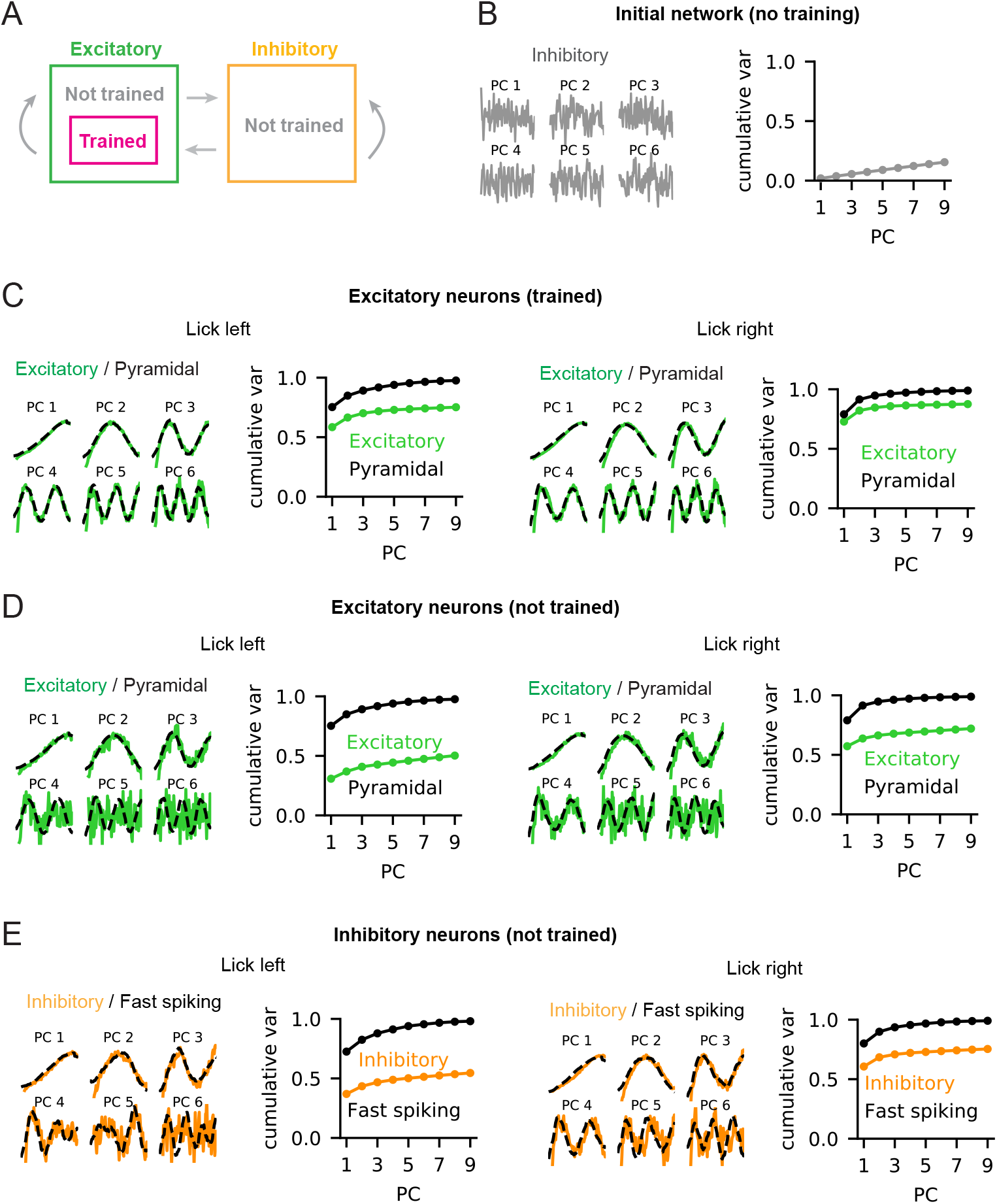
PCA on the PSTHs of the initial network, trained excitatory neurons, untrained excitatory neurons and untrained inhibitory neurons. **(A)** Schematic of the Subset Training, where only 1824 excitatory neurons out of total 2500 excitatory neurons were trained. It is the same trained network presented in Fig. 2. **(B)** The first six PCs of PSTHs of the inhibitory neurons in the initial network before training (left). The cumulative variance explained by the PCs (right). **(C)** PCA on the PSTHs of the trained excitatory model neurons and pyramidal ALM neurons for the lick-left and lick-right trial types. **(D)** PCA on the PSTHs of the untrained excitatory model neurons and pyramidal ALM neurons for the lick-left and lick-right trial types. **(E)** PCA on the PSTHs of the untrained inhibitory model neurons and fast-spiking ALM neurons for the lick-left and lick-right trial types.

**Figure S4:**
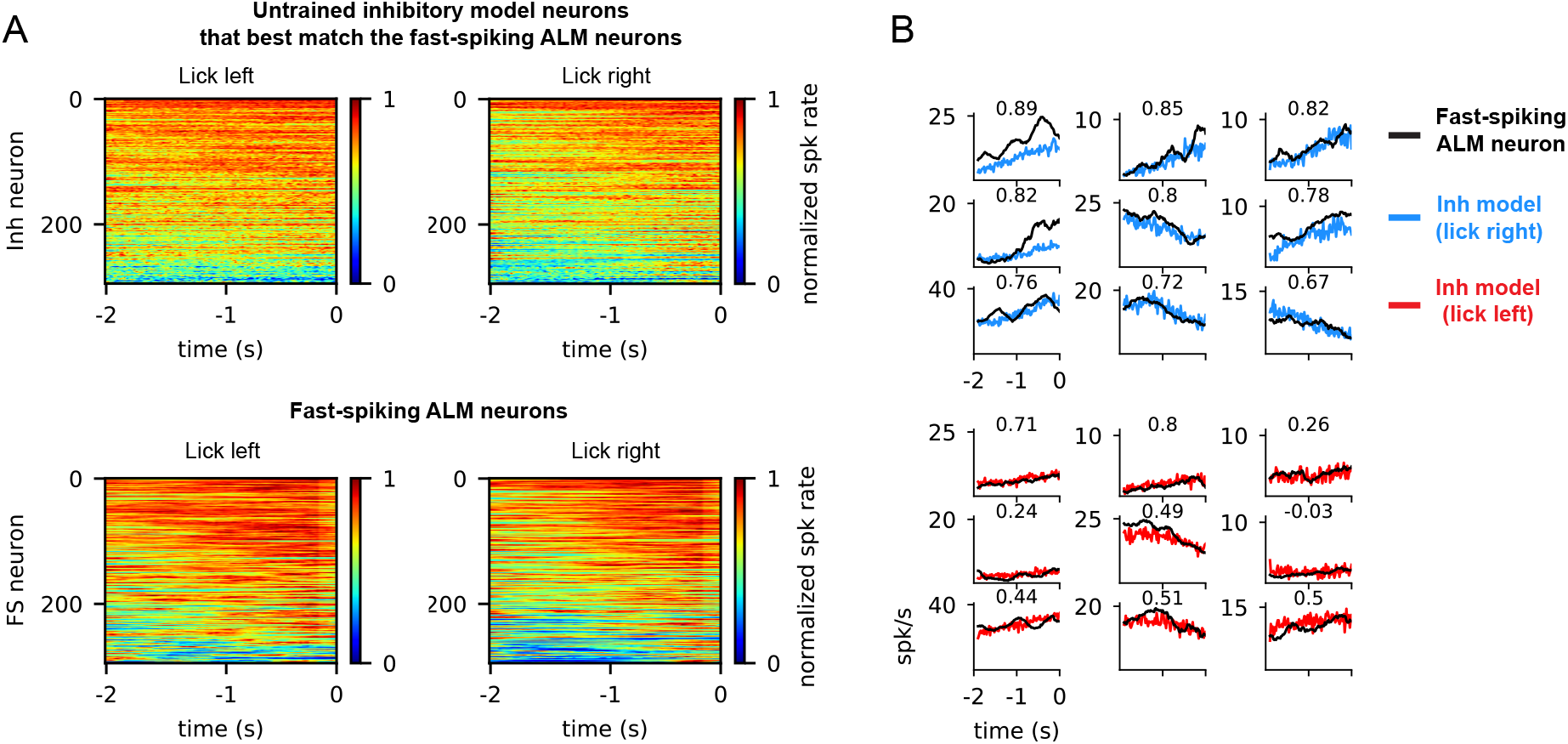
Spiking activity of untrained inhibitory neurons that best match the spiking activity of fast-spiking ALM neurons. **(A)** Normalized PSTHs of the untrained inhibitory neurons (top) selected from the inhibitory population to match the PSTHs of fast-spiking ALM neurons (bottom). Neurons were sorted in decreasing order according to the goodness-of-fit for the lick-right trial. **(B)** Example PSTHs of untrained inhibitory neurons and the matched fast-spiking ALM neurons. Panels in the same position represent the activity of same neurons during the lick-right (top) and lick-left (bottom) trials. The correlation between the matched PSTHs are shown on each panel.

**Figure S5:**
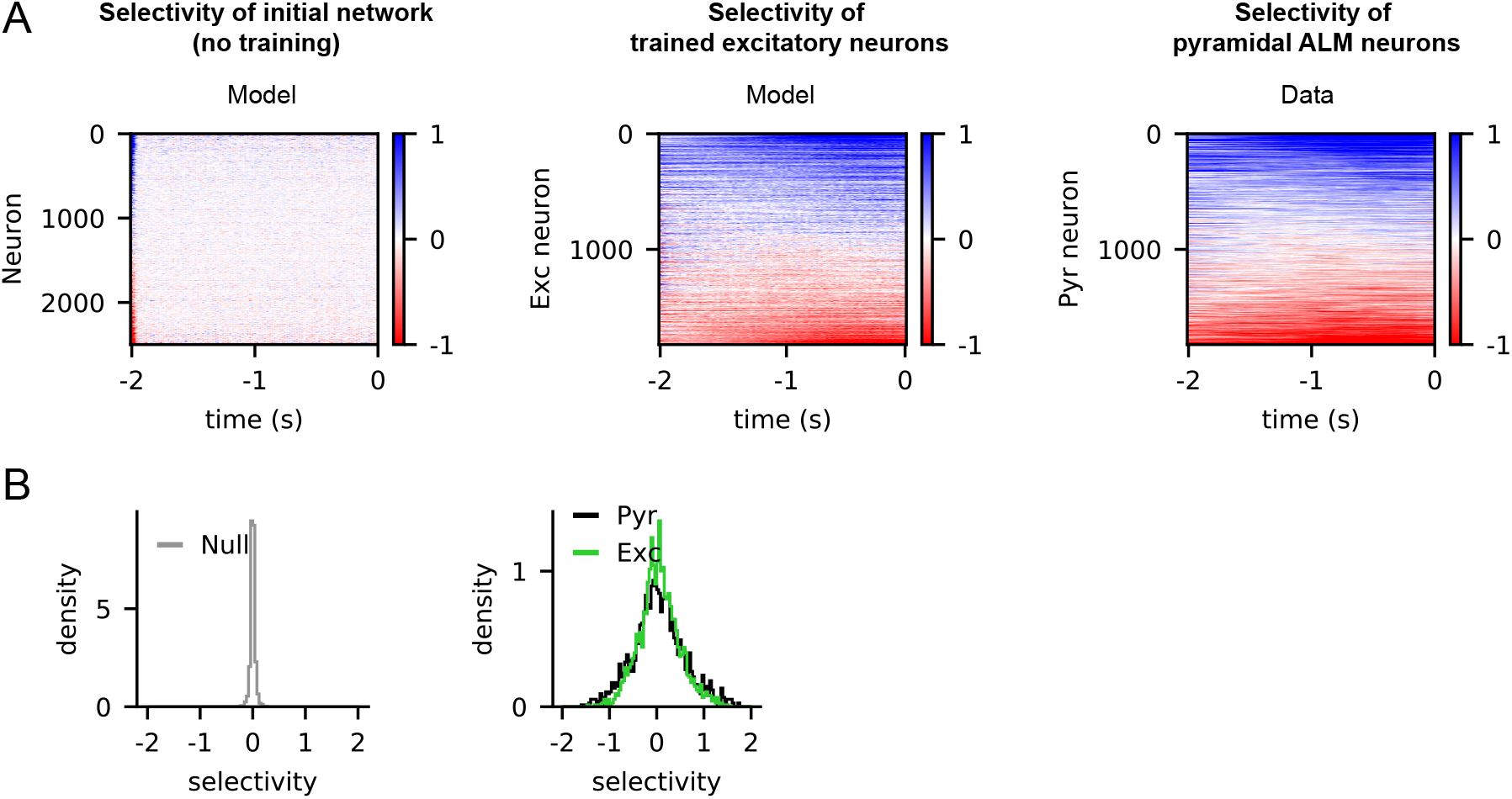
Selectivity of null network neurons, trained excitatory neurons and pyramidal ALM neurons. The null network is the initial balanced network with no training. **(A)** Selectivity of excitatory neurons in the null network (left). Selectivity of excitatory neurons trained to generate the activity of pyramidal ALM neurons (middle). Selectivity of pyramidal ALM neurons (right). **(B)** Distributions of neurons’ choice selectivity in the null network (left), the trained excitatory model neurons and pyramidal ALM neurons (right).

**Figure S6:**
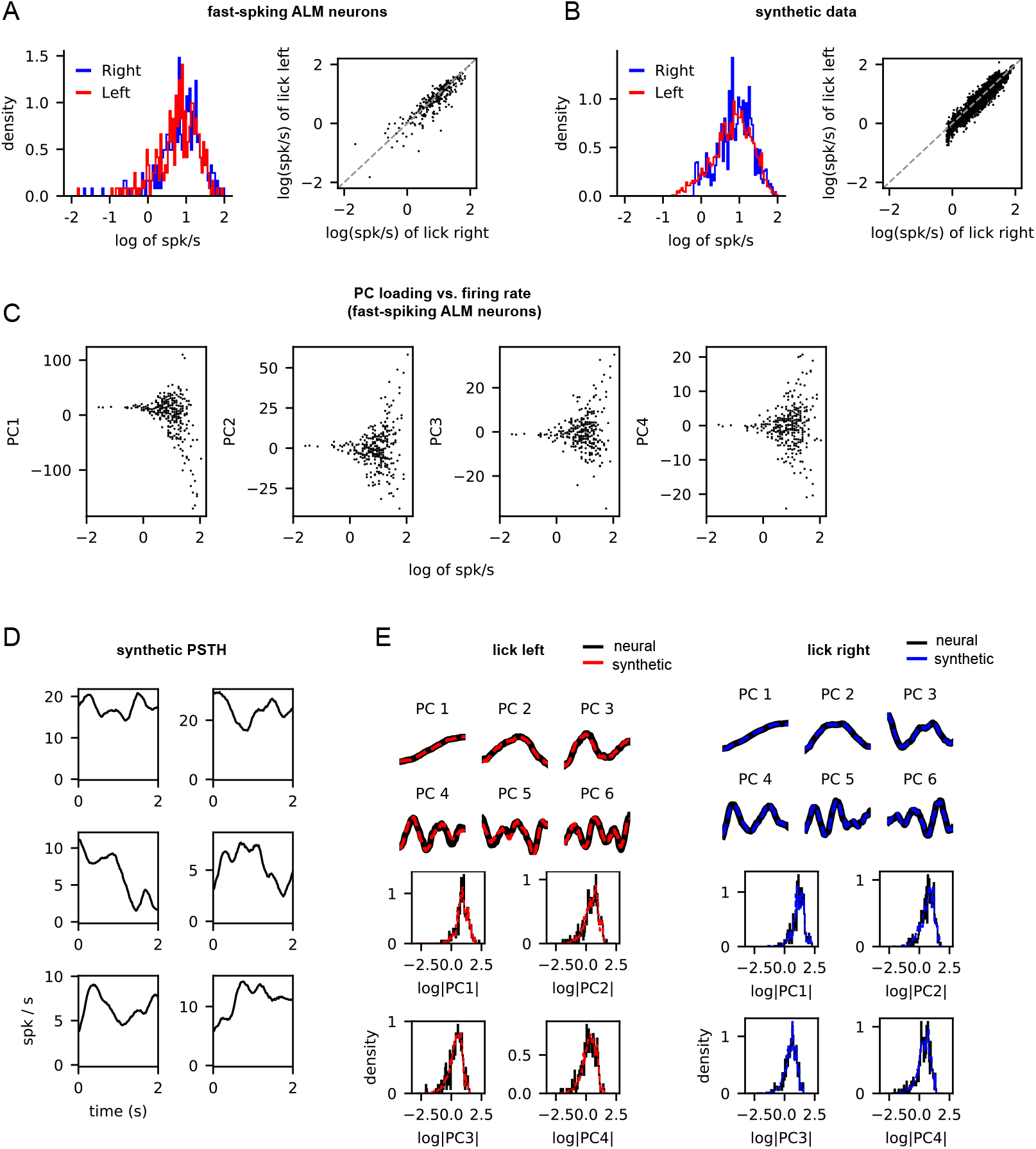
Constructing synthetic neural data. **(A)** Firing rate distribution of fast-spiking ALM neurons for the lick-left and lick-right trials (left). Comparison of neuron’s firing rate for two conditions (right). **(B)** Firing rate distribution of synthetic neurons. The firing rate of a synthetic neuron for the lick-right trial is sampled from ALM neuron’s empirical rate distribution shown in (A, left). To get the firing rate of lick left-trials that has choice selectivity similar to (A, right), we selected fast-spiking ALM neurons whose lick-right firing rate was close to the sampled rate, and added a noise term with mean 0 and variance identical to the empirical variance of chosen ALM neuron’s lick left rates. **(C)** PC loading vs. firing rate of fast-spiking ALM neurons for the lick-right trials. Based on the sampled rates from (B), the PC loadings to each synthetic neuron were bootstrapped from the empirical distribution of fast-spiking ALM neuron’s PC loadings. **(D)** Synthetic PSTHs were constructed by 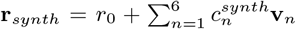 where *r*_0_ is the baseline firing rate from (B), 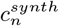 is the *n^th^* PC’s loading from (C) and **v**_*n*_ is the *n^th^* PC of fast-spiking ALM neurons. **(E)** PCA on the PSTHs of fast-spiking ALM neurons and fast-spiking synthetic neurons. The first six PCs (top) and the distributions of PC loadings (bottom) were identical. See also methods for detailed explanation.

**Figure S7:**
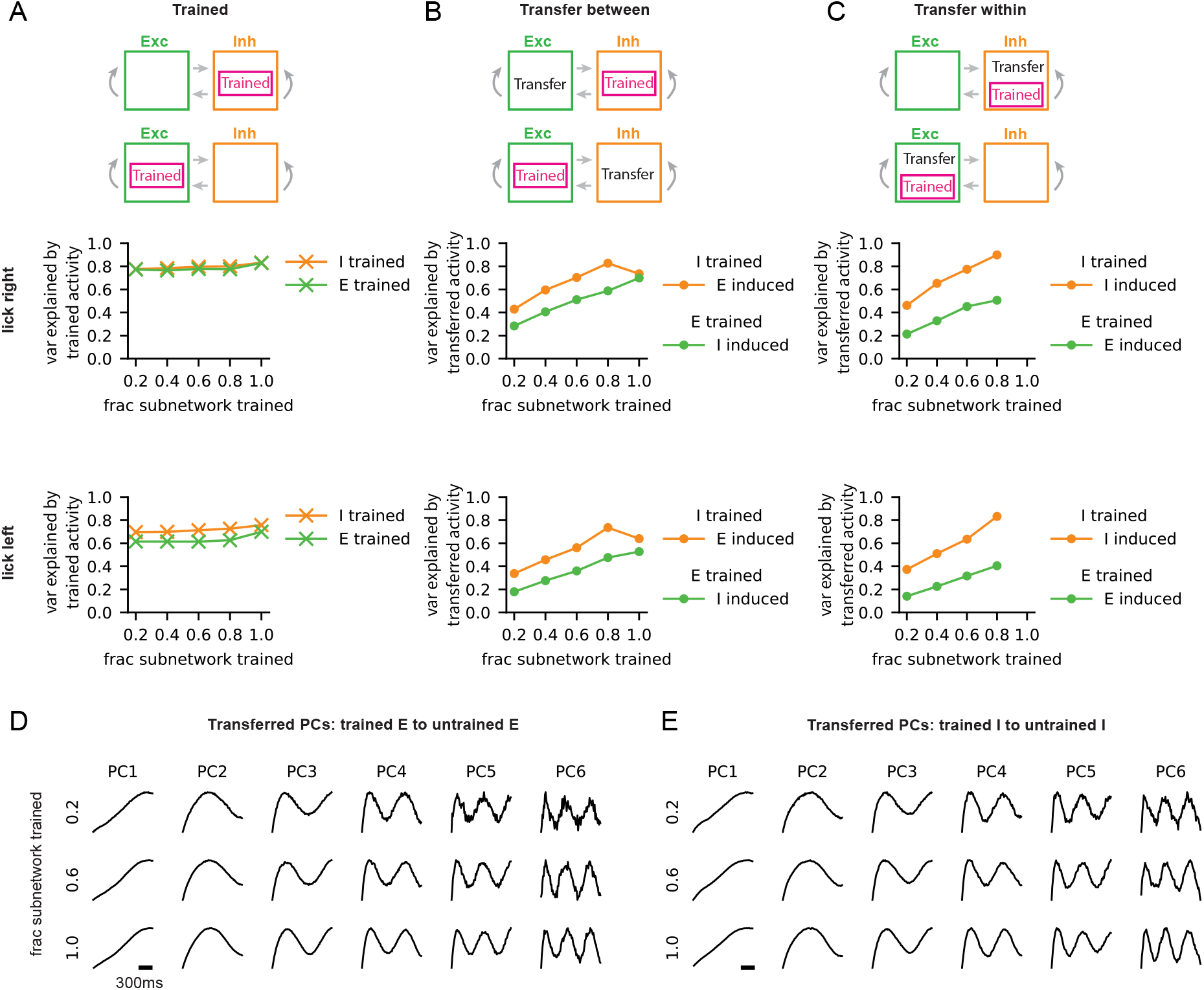
Spreading of trained synthetic neural activity. Extended results from the network trained on synthetic neural data in Fig. 4. A network of size *N* = 30000 was trained using the synthetic neural data. The fraction of trained neurons within the subnetwork was varied, where the fraction equals to 1 if all the neurons within the (excitatory or inhibitory) subnetwork are trained. We considered two training scenarios (top row) where either the excitatory or the inhibitory neurons, but not both, were trained. Explained variance refers to the variance explained by the first six PCs of the PSTHs of neurons that were either trained (panel (A)) or not trained (panels (B) and (C)). **(A)** Variance explained by the subsets of trained neurons in two training scenarios for the lick-right (middle) and lick-left (bottom) trial types. **(B)** Variance explained by the untrained neurons within the subnetwork that was not trained. Two training scenarios for the lick-right (middle) and lick-left (bottom) trial types are shown. **(C)** Variance explained by the untrained neurons within the trained subnetwork. **(D)** PCs of the PSTHs transferred from the trained excitatory neurons within the trained excitatory subnetwork to the untrained excitatory neurons within the same trained excitatory subnetwork. The fraction of trained neurons was varied. This corresponds to the scenarios shown in (C). **(E)** Same as in (D), but now considered the PCs of the PSTHs transferred from the trained inhibitory neurons within the trained inhibitory subnetwork to the untrained inhibitory neurons within the same trained inhibitory subnetwork.

**Figure S8:**
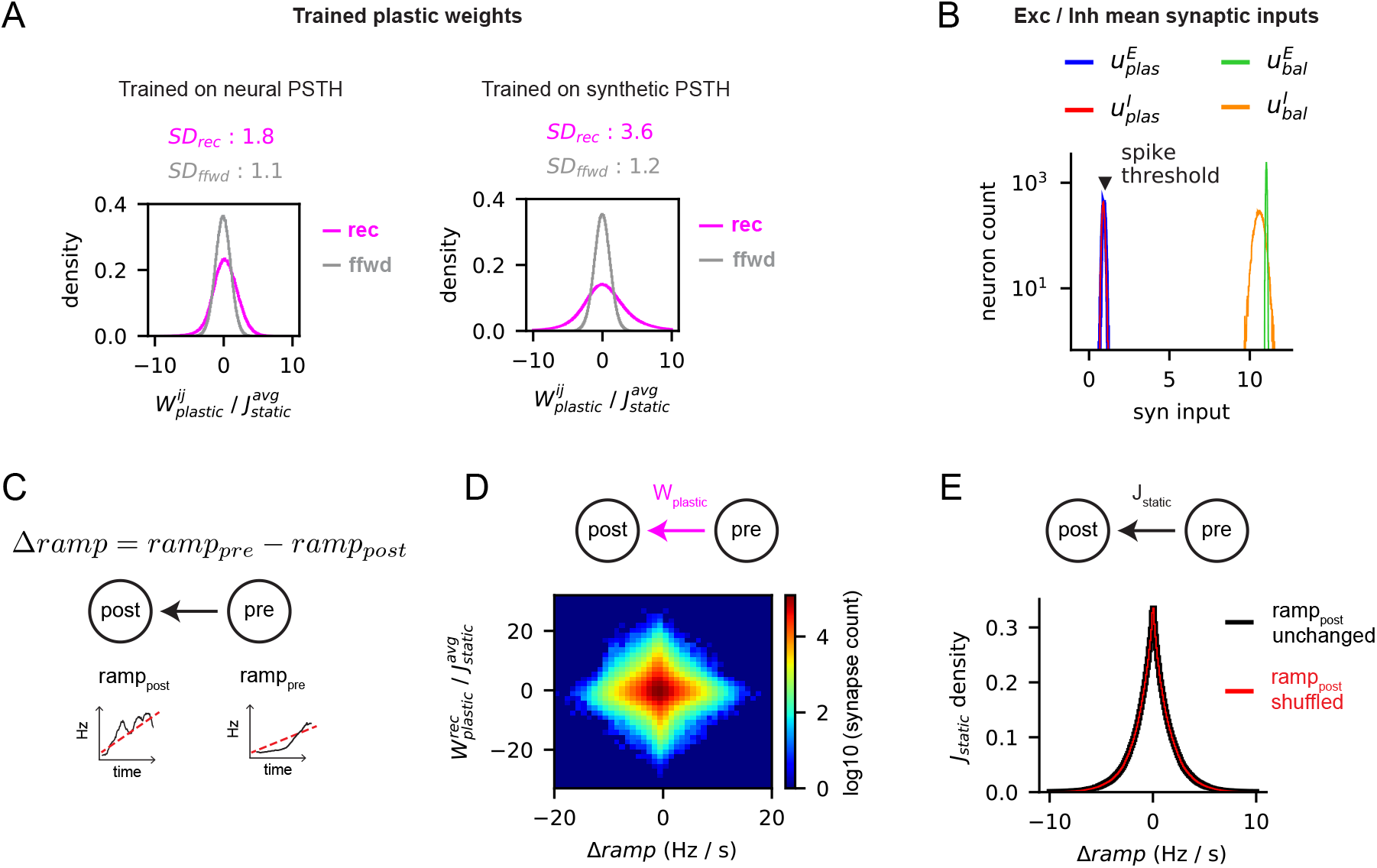
Trained plastic weights are similarly strong as the static weights and correlated to ramping activity. **(A)** Trained plastic weights (recurrent and feedforward), normalized by the average static weights. Left: trained network in Fig. 2 where the 1824 excitatory neurons (out of *N_E_* = 2500) were trained on neural PSTHs. Right: trained network in Fig. 4 where 40% of excitatory neurons (out of *N_E_* = 15000) were trained on synthetic PSTH. Dale’s principle was not respected by the plastic weights in both trained networks. SD shows the standard deviation of the normalized plastic weight distributions. **(B)** Temporal average of excitatory and inhibitory synaptic inputs to trained neurons from the network shown in (A, right). Even though the plastic weights were moderately strong, the excitatory-inhibitory plastic inputs were around spike-threshold and significantly weaker than the balanced inputs. Since the plastic weights did not respect the Dale’s principle, synaptic inputs to a neuron through the positive and negative plastic weights are defined as 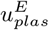 and 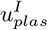, respectively. **(C)** Schematic showing that *ramp_pre_* and *ramp_post_* are the slopes of the linear fit on pre- and post-synaptic neuron’s PSTH, respectively. Δ*ramp* is the difference of the slopes. **(D)** Correlated structure between Δ*ramp* (i.e., difference between the ramping rates of pre- and post-synaptic neurons) and the normalized 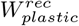 (i.e., strength of the plastic synapse connecting two neurons). Strong plastic weights around Δ*ramp* = 0 gradually decreased as the absolute value of Δ*ramp* increases. **(E)** Unlike the plastic weights shown in (D), the static weights only depended on four connection types (i.e. *E* → *E, E* → *I, I* → *E, I* → *I*). We tested if the ramping activity in untrained neurons might have emerged from preferential inputs from presynaptic neurons sharing similar ramping activity. We evaluated Δ*ramp* between untrained postsynaptic neurons and their presynaptic neurons. The density of static synapses as a function of Δ*ramp* is shown in black. Next, *ramp_post_* was randomly shuffled and Δ*ramp* was re-evaluated. The density of static synapses as a function of the shuffled Δ*ramp* is shown in red. Two distributions were identical, showing the absence of strong correlation between the ramping activity of neurons connected by static synapses.

**Figure S9:**
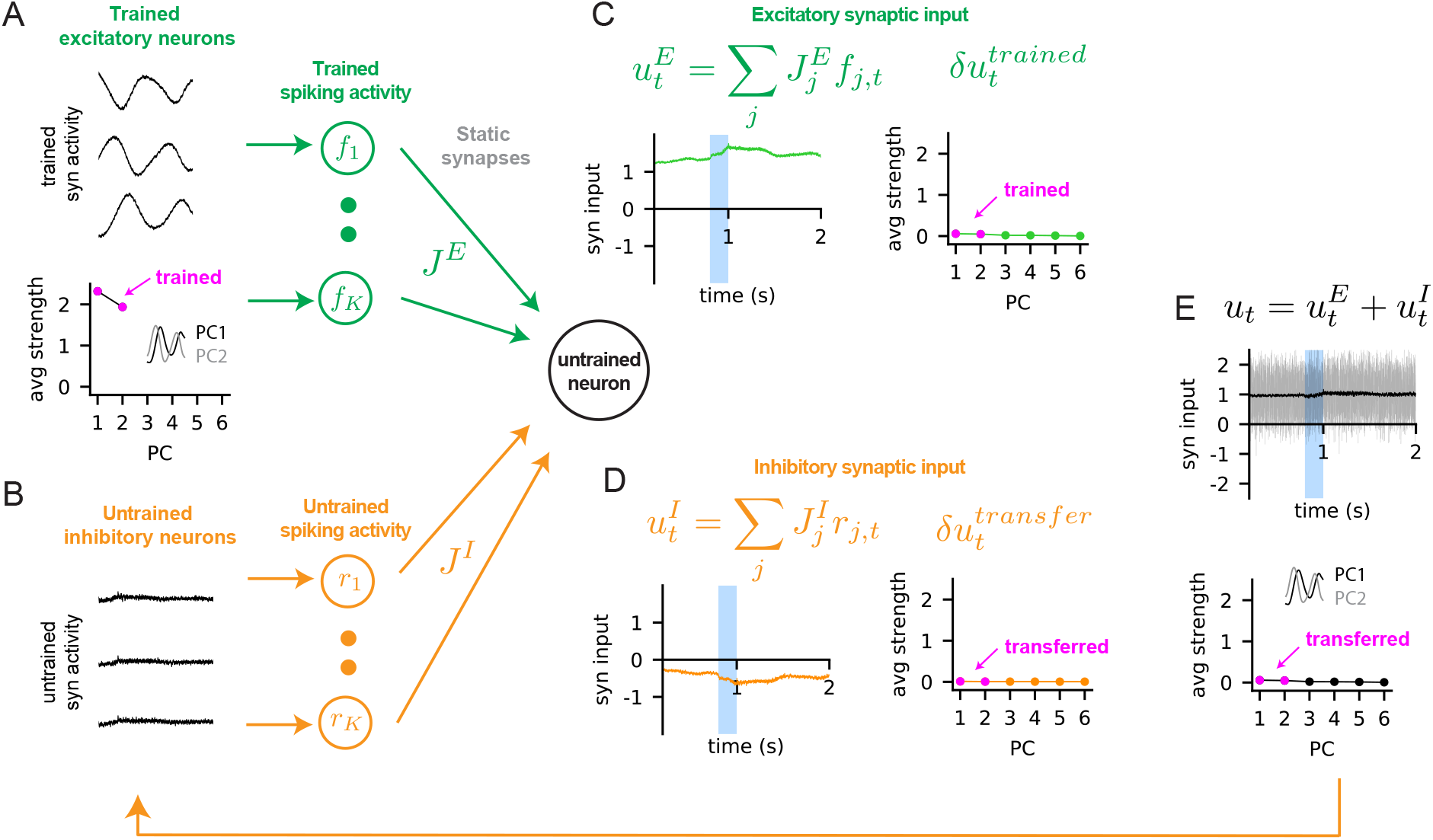
Trained neural activity fails to spread to untrained neurons if the static synapses are weak. To create a weakly connected network, the weights of static synapses were scaled by 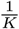, instead of 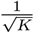. The external inputs were adjusted to be around the spike-threshold such that the excitatory and inhibitory firing rates were around 5Hz and 12Hz, respectively, before training. Gaussian noise with mean 0 and standard deviation 0.5 was injected to the membrane equation to emulate the noisy spiking activity of the balanced network. The excitatory neurons were trained to learn sine waves with frequency 2Hz and random phases, but the inhibitory neurons were not trained. **(A)** The synaptic activities of trained excitatory neurons followed the target sine waves (top). The PCs of the trained excitatory neurons’ synaptic activity (bottom). The Fourier modes of sine waves are highlighted (magenta) and the corresponding PCs (i.e., PC1, PC2) are shown. The loading of each PC on synaptic activity was averaged over all excitatory neurons to obtain the average strength of the PCs. The first two PCs explained close to 99% of the variance. **(B)** Synaptic activity of untrained inhibitory neurons had no temporal structure. **(C)** Aggregate excitatory synaptic input to an untrained inhibitory neuron (left, 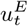). The PCs of the temporal modulation of excitatory input (right, 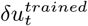) with the corresponding PCs in panel (A) highlighted (magenta). The strength of the trained PCs (i.e., PC1 and PC2) was weak. **(D)** Same as in (C), but the aggregate untrained inhibitory input to the same untrained inhibitory neuron is shown. **(E)** The total synaptic input to the untrained inhibitory neuron (top) with the Gaussian noise shown in the background (gray). The PCs of the total input to untrained inhibitory neurons (bottom) with the transferred PCs highlighted (magenta). Again, the strength of the transferred PCs was weak.

## References

[1] Tsai-Wen Chen, Nuo Li, Kayvon Daie, and Karel Svoboda. A map of anticipatory activity in mouse motor cortex. Neuron, 94(4):866–879, 2017.

[2] William E Allen, Isaac V Kauvar, Michael Z Chen, Ethan B Richman, Samuel J Yang, Ken Chan, Viviana Gradinaru, Benjamin E Deverman, Liqun Luo, and Karl Deisseroth. Global representations of goal-directed behavior in distinct cell types of mouse neocortex. Neuron, 94(4):891–907, 2017.

[3] Hiroshi Makino, Chi Ren, Haixin Liu, An Na Kim, Neehar Kondapaneni, Xin Liu, Duygu Kuzum, and Takaki Komiyama. Transformation of cortex-wide emergent properties during motor learning. Neuron, 94(4):880–890, 2017.

[4] Nicholas A Steinmetz, Peter Zatka-Haas, Matteo Carandini, and Kenneth D Harris. Distributed coding of choice, action and engagement across the mouse brain. Nature, 576(7786):266–273, 2019.

[5] William E Allen, Michael Z Chen, Nandini Pichamoorthy, Rebecca H Tien, Marius Pachitariu, Liqun Luo, and Karl Deisseroth. Thirst regulates motivated behavior through modulation of brainwide neural population dynamics. Science, 364(6437):eaav3932, 2019.

[6] Richard H Roth, Robert H Cudmore, Han L Tan, Ingie Hong, Yong Zhang, and Richard L Huganir. Cortical synaptic ampa receptor plasticity during motor learning. Neuron, 105(5):895–908, 2020.

[7] Simon X Chen, An Na Kim, Andrew J Peters, and Takaki Komiyama. Subtype-specific plasticity of inhibitory circuits in motor cortex during motor learning. Nature neuroscience, 18(8):1109–1115, 2015.

[8] Tonghui Xu, Xinzhu Yu, Andrew J Perlik, Willie F Tobin, Jonathan A Zweig, Kelly Tennant, Theresa Jones, and Yi Zuo. Rapid formation and selective stabilization of synapses for enduring motor memories. Nature, 462(7275):915–919, 2009.

[9] Guang Yang, Feng Pan, and Wen-Biao Gan. Stably maintained dendritic spines are associated with lifelong memories. Nature, 462(7275):920–924, 2009.

[10] Sadegh Nabavi, Rocky Fox, Christophe D Proulx, John Y Lin, Roger Y Tsien, and Roberto Malinow. Engineering a memory with ltd and ltp. Nature, 511(7509):348–352, 2014.

[11] Yann Humeau and Daniel Choquet. The next generation of approaches to investigate the link between synaptic plasticity and learning. Nature neuroscience, 22(10):1536–1543, 2019.

[12] Kiwamu Takemoto, Hiroko Iwanari, Hirobumi Tada, Kumiko Suyama, Akane Sano, Takeharu Nagai, Takao Hamakubo, and Takuya Takahashi. Optical inactivation of synaptic ampa receptors erases fear memory. Nature Biotechnology, 35(1):38–47, 2017.

[13] Akiko Hayashi-Takagi, Sho Yagishita, Mayumi Nakamura, Fukutoshi Shirai, Yi I Wu, Amanda L Loshbaugh, Brian Kuhlman, Klaus M Hahn, and Haruo Kasai. Labelling and optical erasure of synaptic memory traces in the motor cortex. Nature, 525(7569):333–338, 2015.

[14] Wataru Kakegawa, Akira Katoh, Sakae Narumi, Eriko Miura, Junko Motohashi, Akiyo Takahashi, Kazuhisa Kohda, Yugo Fukazawa, Michisuke Yuzaki, and Shinji Matsuda. Optogenetic control of synaptic ampa receptor endocytosis reveals roles of ltd in motor learning. Neuron, 99(5):985–998, 2018.

[15] Henry Markram, Joachim Lübke, Michael Frotscher, and Bert Sakmann. Regulation of synaptic efficacy by coincidence of postsynaptic aps and epsps. Science, 275(5297):213–215, 1997.

[16] Guo-qiang Bi and Mu-ming Poo. Synaptic modifications in cultured hippocampal neurons: dependence on spike timing, synaptic strength, and postsynaptic cell type. Journal of neuroscience, 18(24):10464–10472, 1998.

[17] Shinichiro Tsutsumi and Akiko Hayashi-Takagi. Optical interrogation of multi-scale neuronal plasticity underlying behavioral learning. Current Opinion in Neurobiology, 67:8–15, 2021.

[18] Dimitry Fisher, Itsaso Olasagasti, David W Tank, Emre RF Aksay, and Mark S Goldman. A modeling framework for deriving the structural and functional architecture of a short-term memory microcircuit. Neuron, 79(5):987–1000, 2013.

[19] Kanaka Rajan, Christopher D. Harvey, and David W. Tank. Recurrent Network Models of Sequence Generation and Memory. Neuron, 90(1):128–142, 2016.

[20] Kayvon Daie, Karel Svoboda, and Shaul Druckmann. Targeted photostimulation uncovers circuit motifs supporting short-term memory. Nature Neuroscience, 24(2):259–265, 2021.

[21] Arseny Finkelstein, Lorenzo Fontolan, Michael N Economo, Nuo Li, Sandro Romani, and Karel Svoboda. Attractor dynamics gate cortical information flow during decision-making. Nature Neuroscience, 24(6):843–850, 2021.

[22] Aaron S Andalman, Vanessa M Burns, Matthew Lovett-Barron, Michael Broxton, Ben Poole, Samuel J Yang, Logan Grosenick, Talia N Lerner, Ritchie Chen, Tyler Benster, et al. Neuronal dynamics regulating brain and behavioral state transitions. Cell, 177(4):970–985, 2019.

[23] Carl Van Vreeswijk and Haim Sompolinsky. Chaos in neuronal networks with balanced excitatory and inhibitory activity. Science, 274(5293):1724–1726, 1996.

[24] Nicolas Brunel. Dynamics of sparsely connected networks of excitatory and inhibitory spiking neurons. Journal of computational neuroscience, 8(3):183–208, 2000.

[25] Alfonso Renart, Jaime De La Rocha, Peter Bartho, Liad Hollender, Néstor Parga, Alex Reyes, and Kenneth D Harris. The asynchronous state in cortical circuits. science, 327(5965):587–590, 2010.

[26] Michael N Shadlen and William T Newsome. The variable discharge of cortical neurons: implications for connectivity, computation, and information coding. Journal of neuroscience, 18(10):3870–3896, 1998.

[27] Alex Roxin, Nicolas Brunel, David Hansel, Gianluigi Mongillo, and Carl van Vreeswijk. On the distribution of firing rates in networks of cortical neurons. Journal of Neuroscience, 31(45):16217–16226, 2011.

[28] JUN Tanji and Edward V Evarts. Anticipatory activity of motor cortex neurons in relation to direction of an intended movement. Journal of neurophysiology, 39(5):1062–1068, 1976.

[29] Eberhard E Fetz. Are movement parameters recognizably coded in the activity of single neurons? Behavioral and brain sciences, page 154, 1992.

[30] Mark M Churchland, John P Cunningham, Matthew T Kaufman, Stephen I Ryu, and Krishna V Shenoy. Cortical preparatory activity: representation of movement or first cog in a dynamical machine? Neuron, 68(3):387–400, 2010.

[31] Hidehiko K Inagaki, Miho Inagaki, Sandro Romani, and Karel Svoboda. Low-dimensional and monotonic preparatory activity in mouse anterior lateral motor cortex. Journal of Neuroscience, 38(17):4163–4185, 2018.

[32] Zengcai V Guo, Nuo Li, Daniel Huber, Eran Ophir, Diego Gutnisky, Jonathan T Ting, Guoping Feng, and Karel Svoboda. Flow of cortical activity underlying a tactile decision in mice. Neuron, 81(1):179–194, 2014.

[33] Robert Rosenbaum, Matthew A Smith, Adam Kohn, Jonathan E Rubin, and Brent Doiron. The spatial structure of correlated neuronal variability. Nature neuroscience, 20(1):107–114, 2017.

[34] Ho Ko, Sonja B Hofer, Bruno Pichler, Katherine A Buchanan, P Jesper Sjöström, and Thomas D Mrsic-Flogel. Functional specificity of local synaptic connections in neocortical networks. Nature, 473(7345):87–91, 2011.

[35] David Sussillo and Larry Abbott. Generating Coherent Patterns of Activity from Chaotic Neural Networks. Neuron, 63(4):544–557, 2009.

[36] Wilten Nicola and Claudia Clopath. Supervised learning in spiking neural networks with force training. Nature communications, 8(1):2208, 2017.

[37] Christopher M Kim and Carson C Chow. Learning recurrent dynamics in spiking networks. eLife, 7:e37124, 2018.

[38] Ashok Litwin-Kumar and Brent Doiron. Slow dynamics and high variability in balanced cortical networks with clustered connections. Nature neuroscience, 15(11):1498–1505, 2012.

[39] Hidehiko K Inagaki, Lorenzo Fontolan, Sandro Romani, and Karel Svoboda. Discrete attractor dynamics underlies persistent activity in the frontal cortex. Nature, 566(7743):212–217, 2019.

[40] Carsen Stringer, Marius Pachitariu, Nicholas Steinmetz, Charu Bai Reddy, Matteo Carandini, and Kenneth D Harris. Spontaneous behaviors drive multidimensional, brainwide activity. Science, 364(6437), 2019.

[41] Nuo Li, Tsai-Wen Chen, Zengcai V Guo, Charles R Gerfen, and Karel Svoboda. A motor cortex circuit for motor planning and movement. Nature, 519(7541):51–56, 2015.

[42] Adil G Khan, Jasper Poort, Angus Chadwick, Antonin Blot, Maneesh Sahani, Thomas D Mrsic-Flogel, and Sonja B Hofer. Distinct learning-induced changes in stimulus selectivity and interactions of gabaergic interneuron classes in visual cortex. Nature neuroscience, 21(6):851–859, 2018.

[43] Farzaneh Najafi, Gamaleldin F Elsayed, Robin Cao, Eftychios Pnevmatikakis, Peter E Latham, John P Cunningham, and Anne K Churchland. Excitatory and inhibitory subnetworks are equally selective during decision-making and emerge simultaneously during learning. Neuron, 105(1):165–179, 2020.

[44] Carl van Vreeswijk and Haim Sompolinsky. Chaotic balanced state in a model of cortical circuits. Neural computation, 10(6):1321–1371, 1998.

[45] Rodrigo Laje and Dean V Buonomano. Robust timing and motor patterns by taming chaos in recurrent neural networks. Nature Neuroscience, 16(7):925–933, may 2013.

[46] Dimitri M Kullmann, Alexandre W Moreau, Yamina Bakiri, and Elizabeth Nicholson. Plasticity of inhibition. Neuron, 75(6):951–962, 2012.

[47] Gianluigi Mongillo, Simon Rumpel, and Yonatan Loewenstein. Inhibitory connectivity defines the realm of excitatory plasticity. Nature neuroscience, 21(10):1463–1470, 2018.

[48] Robert Kim and Terrence J Sejnowski. Strong inhibitory signaling underlies stable temporal dynamics and working memory in spiking neural networks. Nature Neuroscience, 24(1):129–139, 2021.

[49] Sonja B Hofer, Ho Ko, Bruno Pichler, Joshua Vogelstein, Hana Ros, Hongkui Zeng, Ed Lein, Nicholas A Lesica, and Thomas D Mrsic-Flogel. Differential connectivity and response dynamics of excitatory and inhibitory neurons in visual cortex. Nature neuroscience, 14(8):1045–1052, 2011.

[50] David Hansel and Carl van Vreeswijk. The mechanism of orientation selectivity in primary visual cortex without a functional map. Journal of Neuroscience, 32(12):4049–4064, 2012.

[51] Cengiz Pehlevan and Haim Sompolinsky. Selectivity and sparseness in randomly connected balanced networks. PloS one, 9(2):e89992, 2014.

[52] Alexandre Mahrach, Guang Chen, Nuo Li, Carl van Vreeswijk, and David Hansel. Mechanisms underlying the response of mouse cortical networks to optogenetic manipulation. Elife, 9:e49967, 2020.

[53] Ben Poole, Subhaneil Lahiri, Maithra Raghu, Jascha Sohl-Dickstein, and Surya Ganguli. Exponential expressivity in deep neural networks through transient chaos. Advances in neural information processing systems, 29, 2016.

[54] Samuel S Schoenholz, Justin Gilmer, Surya Ganguli, and Jascha Sohl-Dickstein. Deep information propagation. arXiv preprint arXiv:1611.01232, 2016.

[55] Haim Sompolinsky, Andrea Crisanti, and Hans-Jurgen Sommers. Chaos in random neural networks. Physical review letters, 61(3):259, 1988.

[56] Ran Darshan and Alexander Rivkind. Learning to represent continuous variables in heterogeneous neural networks. Cell Reports, 39(1):110612, 2022.

[57] Cody Baker, Vicky Zhu, and Robert Rosenbaum. Nonlinear stimulus representations in neural circuits with approximate excitatory-inhibitory balance. PLoS computational biology, 16(9):e1008192, 2020.

[58] Alessandro Ingrosso and LF Abbott. Training dynamically balanced excitatory-inhibitory networks. PloS one, 14(8):e0220547, 2019.

[59] Christopher M. Kim and Carson C. Chow. Training Spiking Neural Networks in the Strong Coupling Regime. Neural Computation, 33(5):1199–1233, 04 2021.

[60] Francesca Mastrogiuseppe and Srdjan Ostojic. Linking connectivity, dynamics, and computations in low-rank recurrent neural networks. Neuron, 99(3):609–623, 2018.

[61] Friedrich Schuessler, Francesca Mastrogiuseppe, Alexis Dubreuil, Srdjan Ostojic, and Omri Barak. The interplay between randomness and structure during learning in rnns. arXiv preprint arXiv:2006.11036, 2020.

[62] Lior Lebovich, Ran Darshan, Yoni Lavi, David Hansel, and Yonatan Loewenstein. Idiosyncratic choice bias naturally emerges from intrinsic stochasticity in neuronal dynamics. Nature Human Behaviour, 3(11):1190–1202, 2019.

[63] Ben Engelhard, Ran Darshan, Nofar Ozeri-Engelhard, Zvi Israel, Uri Werner-Reiss, David Hansel, Hagai Bergman, and Eilon Vaadia. Neuronal activity and learning in local cortical networks are modulated by the action-perception state. bioRxiv, page 537613, 2019.

[64] Yashar Ahmadian and Kenneth D Miller. What is the dynamical regime of cerebral cortex? Neuron, 109(21):3373–3391, 2021.

[65] Misha V Tsodyks, William E Skaggs, Terrence J Sejnowski, and Bruce L McNaughton. Paradoxical effects of external modulation of inhibitory interneurons. Journal of neuroscience, 17(11):4382–4388, 1997.

[66] Alessandro Sanzeni, Bradley Akitake, Hannah C Goldbach, Caitlin E Leedy, Nicolas Brunel, and Mark H Histed. Inhibition stabilization is a widespread property of cortical networks. Elife, 9:e54875, 2020.

[67] Sadra Sadeh and Claudia Clopath. Patterned perturbation of inhibition can reveal the dynamical structure of neural processing. Elife, 9:e52757, 2020.

[68] Agostina Palmigiano, Francesco Fumarola, Daniel P Mossing, Nataliya Kraynyukova, Hillel Adesnik, and Kenneth D Miller. Structure and variability of optogenetic responses identify the operating regime of cortex. bioRxiv, 2020.

[69] Ran Darshan, William E Wood, Susan Peters, Arthur Leblois, and David Hansel. A canonical neural mechanism for behavioral variability. Nature communications, 8(1):1–13, 2017.

[70] Ran Darshan, Carl Van Vreeswijk, and David Hansel. Strength of correlations in strongly recurrent neuronal networks. Physical Review X, 8(3):031072, 2018.

[71] Henry Clavering Tuckwell. Introduction to theoretical neurobiology: linear cable theory and dendritic structure, volume 1. Cambridge University Press, 1988.

